# Genomic signatures of strawberry domestication and breeding

**DOI:** 10.1101/2023.07.12.548723

**Authors:** Zhen Fan, Vance M. Whitaker

**Author notes:** Corresponding author: Vance M. Whitaker < >.

## Abstract

Cultivated strawberry (*Fragaria × ananassa*) has a brief history of less than 300 years, beginning with the hybridization of octoploids *F. chiloensis* and F. *virginiana*. Here we explored the genomic signatures of this history using whole-genome sequences of 289 wild, heirloom and modern varieties. Four non-admixed wild octoploid populations were identified, with recurrent introgression among the sympatric populations. The proportion of *F. virginiana* ancestry increased by 20% in modern varieties over initial hybrids, and the proportion of *F. chiloensis* subsp. pacifica rose from 0 to 3.4%. Effective population size rapidly declined during early breeding. Meanwhile, divergent selection for distinct environments reshaped wild allelic origins in 21 out 28 chromosomes. Despite 20 breeding cycles since the initial hybridization, more than half of loci underlying yield and fruit size are still not under selection. These insights add clarity to the domestication and breeding history of what is now the most widely cultivated fruit in the world.

## Introduction

Cultivated strawberry, (*Fragaria × ananassa* Duchesne ex Rozier; 2n = 8× = 56), arose as a homoploid interspecific hybrid between the allo-octoploid species *F. virginiana* and *F. chiloensis*^1^. The hybridization of these New World species occurred in the 17^th^ century in Europe, making strawberry one of the world’s most recently domesticated crops. Initial hybrids were further crossed with additional wild octoploid accessions and have since been bred in a wide variety of climates and cultural systems^1,2^. Today the worldwide economic value of strawberries is more than 22 billion USD per year^3^.

It is widely accepted that there are four extant wild octoploid strawberry species, *Fragaria chiloensis, F. virginiana*, and the natural hybrids *F. × ananassa* subsp. *cuneifolia* and *F. iturupensis* (only found on Iturup Island, Japan). Both *F. chiloensis* and *F. virginiana* are endemic to the Americas and span a wide range of ecological amplitude. Few studies have addressed the taxonomy of the octoploid species. The recent revised taxonomy of the genus recognized four subspecies in *F. virginiana* and five subspecies in *F. chiloensis* based on geographic distribution and morphological characteristics^4^. In one study, morphological traits discriminated three populations in *F. virginiana* (subsp. platypetala, subsp. glauca and subsp. *virginiana*), while marker data only supported two distinct groups: *F. virginiana* subsp. *platypetala* and the rest^5^. Within *F. chiloensis*, SNP array data supported distinct North American and South American populations^6^. Since the wild octoploid species are interfertile, introgression between two species and a hybrid zone were proposed^6,7^. Higher confidence in the classification of species and subspecies and the extent of introgression among them are needed to better define their contributions to cultivated strawberry.

Hybridization plays an important role in crop domestication, especially in clonally propagated crops^8^. Apple (*Malus domestica*) underwent hybridization with *Malus sylvestris* during its expansion along the Silk Route^9,10^. Another tree crop, citrus, showed extensive admixture for many species in the genus^11,12^. Hybridization facilitates crop domestication in two aspects: it combines the best characteristics of both parental species and increases genetic diversity amenable for human selection. In citrus, pummelo admixture into mandarins, oranges, and grapefruits was associated with larger fruit size and higher acidity^11^. In grape, pervasive hybridization of local wild relatives with Western European grapevine varieties facilitated environmental adaptations^13^.

Although *F. chiloensis* has sizable fruit and was cultivated by native peoples in Chile and Ecuador for perhaps 1000 years^14^, its lack of cold hardiness and heat tolerance prevented its cultivation in North America (NA) and other parts of the world. Hybridization with *F. virginiana* infused hardiness that adapted *F. × ananassa* to diverse climates, but the transition from native species to modern varieties was lengthy, involving migration, further wild introgressions, and human selection^1,15^. Antoine Nicolas Duchesne, a French horticulturist was the first to recognize that *F. × ananassa* was an interspecific hybrid between *F. chiloensis* and F. *virginiana*^1^. The first generation of *F. × ananassa* varieties developed in NA, which included ‘Hovey’ and ‘Rose Phoenix’ were predicted to be hybrids of European cultivars with native subspecies^1^, although the taxonomic origin of their parents remains a mystery^15^. Since the early 1900s, private growers and public institutions, which included the University of California, Davis (UCD) and the University of Florida (UF) established systematic breeding of *F. × ananassa*. Due to improved domesticated traits like perpetual flowering and adaptation to annualized culture, California cultivars spread throughout the world^15–17^. The UF strawberry program began in 1948, focusing on winter production in a subtropical environment, spreading varieties to other winter and spring production regions^16,18,19^.

The domestication of strawberry has been rapid compared to most seed crops and even other perennial fruit crops which occurred over thousands of years^8,26^. Yet the molecular footprint of this period was not accessible until the recent publication of high-quality genomes^20,21^. Sub-genome-specific DNA variation can now be accurately identified via mapping resequencing data to reference genomes^22^. Hardigan et al.^23^ and Pincot et al.^15^ conducted companion studies, using pedigree records and whole-genome sequencing data of 145 octoploids to examine breeding progress. Contrary to previous assumptions of low genetic diversity^24,25^, domesticated populations maintained over 70% of total alleles from wild populations^23^, possibly benefitted by recurrent hybridization^15^. A higher percentage of the *F. × ananassa* genome was more closely related to *F. virginiana* than *F. chiloensis*^23^. Recurrent selection has left its mark in the genomes of modern varieties, including large linkage decay (400kb) and selective sweeps in both early and modern phases of domestication^23^. A significant GWAS signal for fruit firmness overlapped with early-phase domestication sweeps.

Our objectives of the present study were to examine speciation and subsequent introgressions in wild octoploid populations and to explore remaining questions around cultivated strawberry domestication. These questions include, what proportions of modern genomes are attributable to different wild subspecies; how do wild allelic origins vary by chromosome; how did population size and diversity fluctuate during domestication and breeding; what loci conferred increases in fruit size and yield; and which of these loci are under selection? This was accomplished using DNA variants from whole genome sequencing of a diverse panel of 289 cultivars and wild octoploid progenitors.

## Results

Our WGS study of octoploid strawberry was conducted with 289 re-sequenced octoploid strawberry samples. The newly constructed octoploid strawberry variant map includes a total of 94.94 M SNPs and 15.31 M InDels. Among them, 0.82 M are frameshift InDels, and 5.24 M are nonsynonymous SNVs (Table S3).

### Recurrent natural hybridization between *F. virginiana* and *F. chiloensis*

Included in the 289 re-sequenced octoploid strawberries were 102 wild individuals, from which classification of species and subspecies could be confidently inferred based on genome-wide DNA variants. Both PCA and admixture analysis (K=3) separated cultivated strawberry (*F. × ananassa*), *F. virginiana* and *F. chiloensis* (Fig. 1A & B). Spontaneous hybridization between *F. virginiana* and *F. chiloensis* was detected for the four samples collected at Tucquala Lake, WA, which were previously identified as *F. virginiana* subsp. platypetala^26^. Their ancestry proportions were distinct from samples of *F. × ananassa* subsp. cuneifolia (Fig. 1A), indicating they belonged to a different admixture population. Most *F. virginiana* subsp. *glauca* samples (7 out of 8) were admixtures of two other subspecies in F. *virginiana*. Meanwhile, *F. chiloensis* f. chiloensis (n = 5), *F. chiloensis* subsp. *lucida* (n = 3) and *F. virginiana* subsp. grayana (n = 2) did not form their own clusters in both NJ tree and admixture analyses *with* K = 7 or higher, but nested within *F. chiloensis* f. patagonica (n = 8), *F*. chiloensis subsp. pacifica (n = 27), and *F. virginiana* subsp. *virginiana* (n = 29), respectively.

**Figure 1.**
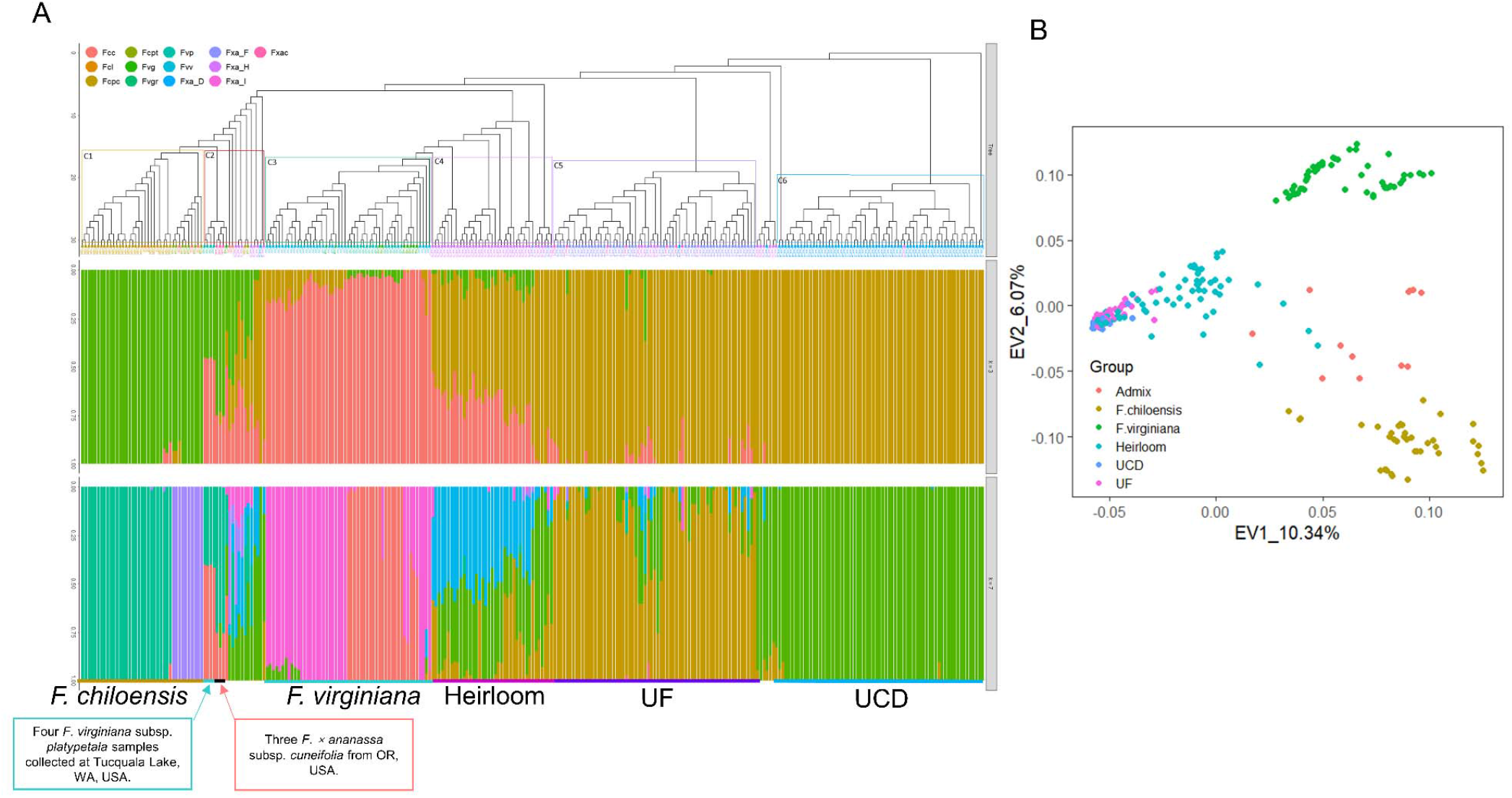
(A) Neighbor-joining tree (top) and admixture analysis of 289 octoploid strawberry individuals (middle and bottom). For admixture analysis, K values of 3 (middle) and 7 (bottom) are plotted. Tips are labeled and colored according to sample taxa or groups. Fcc: *F. chiloensis* f. chiloensis, Fcl: *F. chiloensis* subsp. lucida, Fcpc: *F. chiloensis* subsp. *pacifica*, Fcpt: *F. chiloensis* f. *patagonica*, Fvg: *F. virginiana* subsp. glauca, Fvgr: *F. virginiana* subsp. grayana, Fvp: *F. virginiana* subsp. platypetala, Fvv: *F. virginiana* subsp. *virginiana*, Fxa_D: University of California, Davis accessions (*F. × ananassa*), Fxa_F: University of Florida accessions (*F. × ananassa*), Fxa_H: heirloom varieties (*F. × ananassa*), Fxa_I: recent introgressed lines (*F. × ananassa*), Fxac: *F. × ananassa* ssp. cuneifolia. (B) PCA plot using genetic markers. Variance explained by the top two eigenvectors are included in axes labels. Individual dots are colored by their group.

A species tree built using 1,866,574 SNPs in subgenome A confirmed the monophyletic origin of octoploid strawberry ^27,28^ and F. vesca subsp. bracteata as the diploid donor of subgenome A^20,27,28^ (Fig. 2A). The divergence time between *F. virginiana* and *F. chiloensis* was estimated to be 322 kya (CI: [216, 358]) (Fig. 2A), consistent with a previous estimation using plastid DNA^29^, while the divergence of subspecies was dated to 126 (CI: [52, 138]) and 164 (CI: [146, 216]) kya respectively for subspecies within *F. virginiana* and *F. chiloensis*. In addition to recent wild hybrids (Fig. 1A), a signal for more ancient introgression (f-branch = 0.0923) was found between sympatric *F. chiloensis* subsp. pacifica and *F. virginiana* subsp. platypetala (Fig. 2B). The best Dadi model^30^ also predicted migration between the two species and exponential growth of the population size after divergence (Fig. S2, Table S4). These results demonstrate that spontaneous hybridization between sympatric octoploid Fragaria species was recurrent. However, geographic isolation prevented gene flow into or from *F. chiloensis* f. patagonica (Fig. 2B). Although positive D and *f*4-ratios were observed between diploid and octoploid species (Fig. 2B), a similar level of introgression was revealed for its diploid ancestral species, *F. vesca subp. bracteata* from two east Asian species. Therefore, the observed introgression among interploid species likely happened prior to polyploidization.

**Figure 2.**
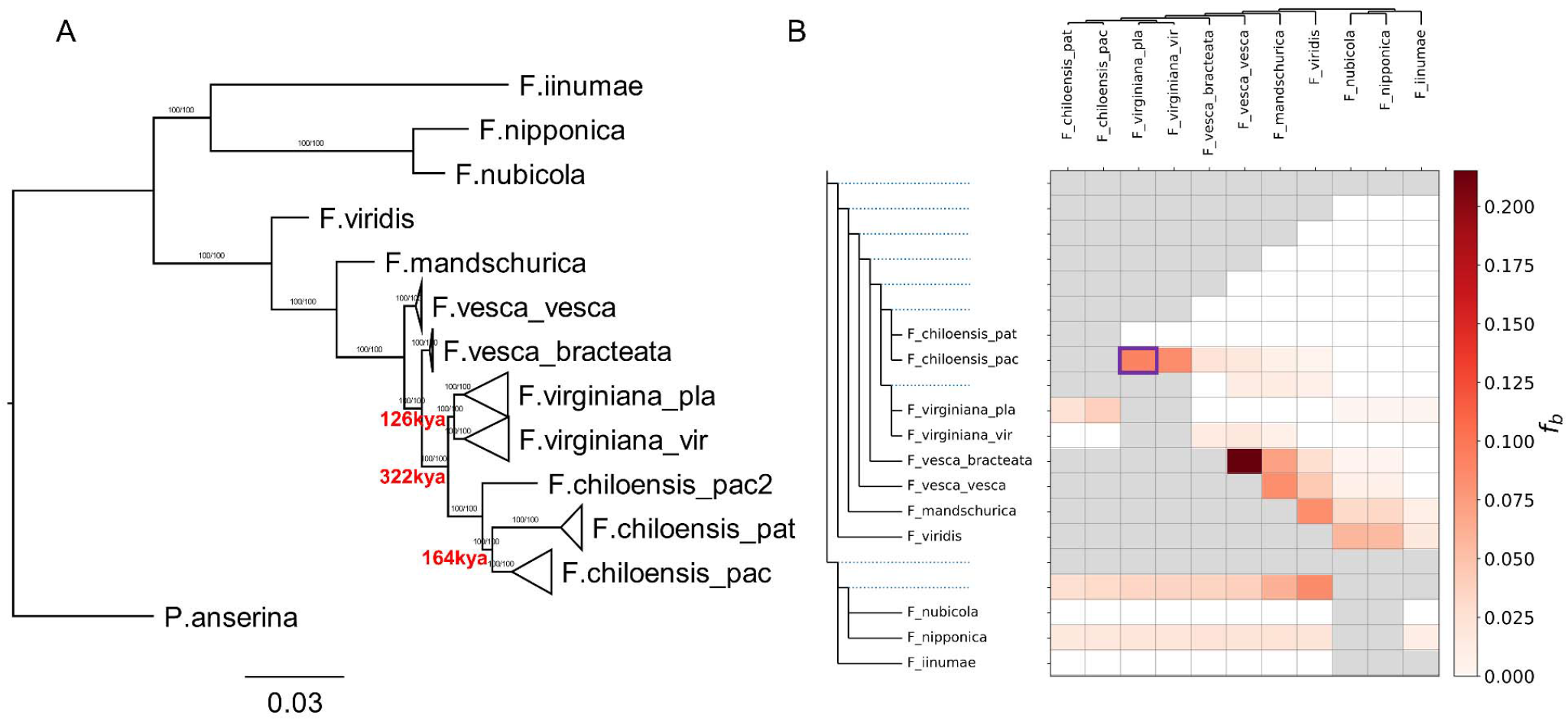
(A) A maximum likelihood species tree using 9,822,133 SNPs on subgenome A (*Fragaria vesca-like*). Branches are labeled with ultrafast bootstrapping value and SH-like approximate likelihood ratio. Divergence time for octoploid species was estimated using SMC++. (B) Heatmap showing F-branch values for different pairs of species. Introgression between octoploid species is indicated by positive F-branch values, one of which is highlighted in a purple box.

### Domestication of *F*. × *ananassa*

Based on PCA and admixture analysis, early breeding accessions (heirlooms) had large proportions of wild genetics. Wild genetic diversity was gradually reduced during breeding (Fig. 1A). However, neither analysis resolved subspecies composition in *F. × ananassa*. Therefore, RFMix^31^ was used to infer local ancestry for all *F. × ananassa* samples. Based on the results from admixture analysis (Fig. 1A), wild species were grouped into four ancestral populations: Fcc/Fcpt (representing *F. chiloensis* subsp. *chiloensis*) including both *F. chiloensis f. chiloensis* and *F. chiloensis f. patagonica* samples; Fcpc/Fcl including both *F. chiloensis* subsp. *pacifica* and *F. chiloensis* subsp. *lucida* samples; Fvp representing *F. virginiana* subsp. *platypetala*; and Fvv/Fvgr including both *F. virginiana* subsp. *virginiana* and *F. virginiana* subsp. *grayana* samples. The results showed that three early varieties from Europe had close to no genetic inheritance from Fcpc/Fcl (Fig. 3B & 3C, Proportion_Fcpc/Fcl_ < 0.01). In contrast almost all post-1900 *F. × ananassa* varieties had a higher proportion of Fcpc/Fcl ancestry (mean = 0.052, 0.032, 0.036 for heirloom, UCD, UF, respectively), and an early California variety, ‘Ettersburg121’, released in 1908^1,17^ was confirmed as a hybrid with Fcpc/Fcl (Fig. 3C). Thus the introgression of Fcpc/Fcl happened after *F. × ananassa* migrated to North America (NA), and its genomic representation has been stationary throughout breeding. Similarly, a higher percentage of Fvv/Fvgr ancestry was observed in varieties developed after the migration to NA (Fig. 3B), in line with previous analyses based on genetic distance^23^. On the other hand, no *F. × ananassa* shared ancestry from Fvp (Fig. 3B). Based on this evidence and previous literature^1,15,17,23^, a detailed domestication route for octoploid strawberry can be visualized according to results from the admixture analysis (Fig. 3A). After French colonel Frézier brought *F. chiloensis* subsp. *chiloensis* plants from Concepcion, Chile to France^1^, initial hybridization between *F. chiloensis* subsp. *chiloensis* and *F. virginiana* subsp. *virginiana* happened around the 1750s in Europe^1^. The human-selected hybrid, *F. × ananassa*, migrated back to the NA in the early 1800s^1^. During 1800s, the pioneer North American breeders crossed early European cultivars with local wild species^1,17^, including *F. chiloensis* subsp. *pacifica* and *F. virginiana* subsp. *virginiana*. Since the mid-1900s, the genome-wide ancestral composition has remained relatively unchanged in varieties across the globe.

**Figure 3.**
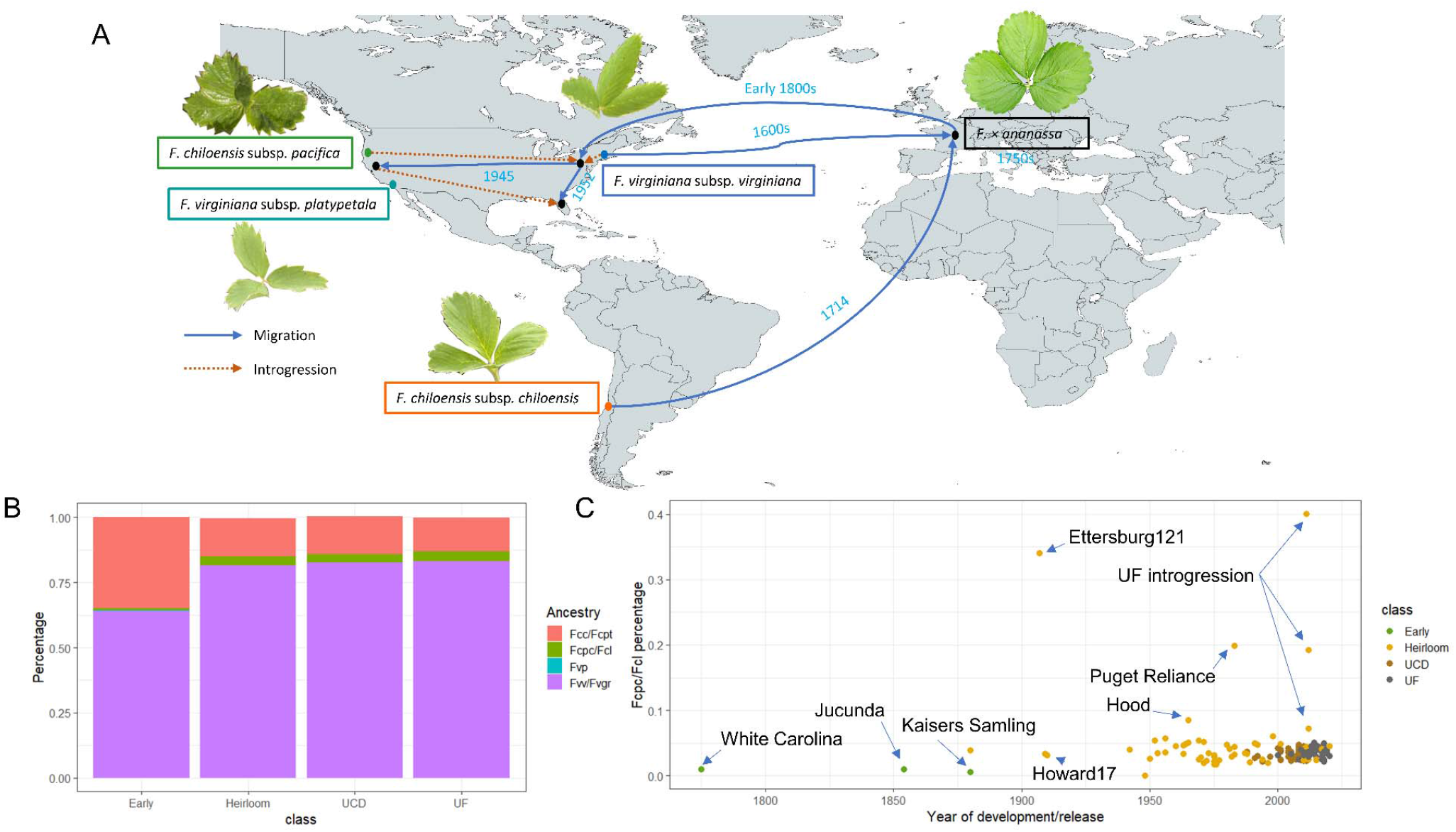
(A) A domestication route for the octoploid strawberry in the Europe and Americas. Dashed lines indicate species/breeding materials introgressed during systematic breeding. (B) The ancestral inference across breeding populations. Early includes three European cultivars released before 1900s, shown in red in figure C. Due to lack of separation in PCA and admixture analyses, *F. chiloensis* f. chiloensis/*F. chiloensis* f. patagonica (Fcc/Fcpt), *F. chiloensis* subsp. *pacifica*/*F. chiloensis subsp. lucida* (Fcpc/Fcl), *F. virginiana* subsp. *virginiana*/*F. virginiana* subsp. *grayana* (Fvv/Fvgr) were respectively combined. Fvp is *F. virginiana* subsp. *platypetala*. (C) The percentage of *F. chiloensis* subsp. *chiloensis* (*F. chiloensis* f. *chiloensis*/*F. chiloensis* f. *patagonica*) ancestry in domesticated varieties.

### Modern breeding reshaped the genome of *F*. × *ananassa*

Since the early 19^th^ century, *F. × ananassa* was continuously bred for California coastal Mediterranean and Florida subtropical climates, among many others. Varieties developed by UCD and UF with improved yield and fruit quality promoted global cultivation of *F. × ananassa*, as the annualized raised-bed culture in these regions gradually became the dominant production method worldwide. Although a similar genome-wide ancestry composition was shared between the two populations (Fig. 3B), significant differences in wild allelic origins were found in 21 out of 28 chromosomes (Wilcoxon signed-rank test, pbonferroni < 0.01), including larger than 10% differences in chromosomes 2A, 4B, 4C, 5B and 7D (Fig. 4A, Table S5), implying progenitor species-preferential selection for distinct production environments. Comparisons among 17 selected founders of the UF population revealed large variation in ancestral species composition among early breeding parents for those five chromosomes (Fig. S3). Although considerable variation in wild allelic origins was preserved in recent breeding lines across some chromosomes by both programs, chromosomes 1B, 3A, 5D and 6B exhibit low variation in the UF population (Fig. S4) and have near zero variation in the UCD population (Fig. S5, Table S5).

**Figure 4.**
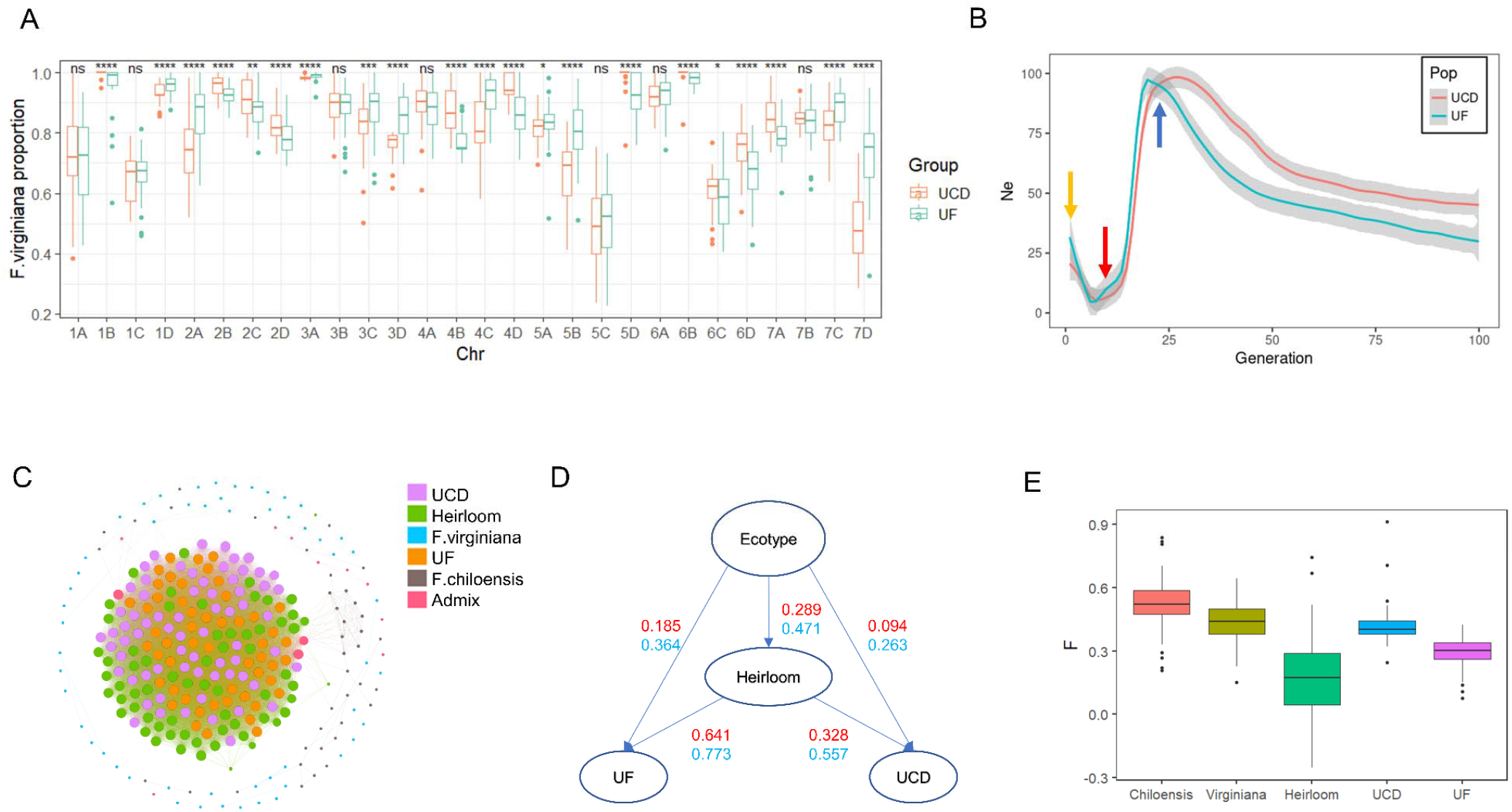
(A) Chromosomal view of fractions of Fragaria *virginiana* ancestry in UCD (orange) and UF (green) populations. Asterisks represent significances for individual chromosomes. (B) Effective population size trajectories for both UCD (orange) and UF populations (green). Three arrows point to respective signs of a recent expansion (yellow), a strong bottleneck (red) and the initial hybridization (blue). (C) Identical by descent network of all sampled individuals. Dots present individuals colored based on their population. The size of a dot is proportional to its weighted IBD which is calculated as the sum of its IBD relationships. (D) Genetic diversity change during domestication. The numbers in red represent the retained proportions of rare alleles (frequencies < 0.2), and numbers in blue represent the retained proportions of total alleles. (E) Inbreeding coefficients (F) across different populations.

Demographic history analysis of both populations reflected the initial hybridization of the two octoploid species, reflected in an effective population size peak (Ne = 86) about 20 generation ago, followed by a strong bottleneck (Ne = 5) around eight generations ago and a subsequent expansion in the last four generations (Fig. 4B, Fig. S6). Current effective population sizes were 18 and 27 for the UCD and UF populations respectively. A distinct, single pulse model32 estimated the admixture time to be 8.39 ± 1.77 and 11.51 ± 1.12 generations for UCD and UF populations respectively (Fig. S7). Within the populations, there was an extensive level of relatedness (Fig. 4C, Table 1). A striking 75.15% of the UCD accessions exhibit second degree or higher relatedness (IBD>0.25), and all the rest showed third to second degree relatedness (0.125<IBD<0.25), compared to 30.62% of higher than second degree relatedness among the UF modern accessions (Table 1). UCD varieties released from 1950 to 1990 were found to be the major contributors to both breeding programs (Fig. S8). In contrast, among the heirloom varieties, only 2.36% individuals shared IBD higher than 0.25. The UCD population was also characterized by longer runs of homozygosity within an individual (Fig. S9). Compared to ecotypes (π = 0.00845) and heirlooms (π = 0.00703), recent breeding populations (πUCD = 0.00428, πUF = 0.00512) showed significantly lower genetic diversity (Fig. S10). The π estimate of ecotype was much higher than a previous estimate23, most likely due to a larger sample collection. Rare alleles with frequencies lower than 0.2 in ecotypes were continuously removed during early domestication and diversification in the modern era (Fig. 4D, Fig. S11). Approximately 18.5% and 9.4% of rare alleles flowed into UF and UCD populations from ecotypes (Fig. 4D). Small numbers of founders and extensive crosses among closely related accessions were likely the main drivers for the bottleneck and decreased diversity. However, the inbreeding coefficients across breeding populations were still lower than wild progenitor populations (Fig. 4E), which can be attributed to the initial interspecific hybridization and a short domestication history.

**Table 1.** IBD relationships in breeding populations

| IBD relationship | Second degree or higher<br>(0.25 < r) | Third to second degree<br>(0.125 < r < 0.25) |
| --- | --- | --- |
| UCD elite | 75.15% (1712/2278) | 24.85% (566/2278) |
| UF elite | 30.62% (406/1326) | 69.38% (920/1326) |
| Heirloom | 2.36% (49/2080) | 38.27% (796/2080) |

### Parallel and divergent selections for different climates/regions

There were 210 selective sweeps associated with the early domestication transition (prior to 1980), covering 22.4% of the genome (Fig. S12, Data S1), and 182 and 172 selective sweeps covering 21.7% of the genome for both the UCD and UF populations (Fig. S13). The proportions of the genome under selection for both the early phase of domestication and the UCD population were larger than previous estimates^23^ and also larger than crops with longer domestication histories^33,34^. These differences were due to a more lenient threshold (5% of the highest selection coefficients) and a merged window of 400kb based on the previous LD decay estimate for the UCD population^23^. Overlapping three scans allowed assignment of selective sweeps into three classes: early selection (present in all three scans), parallel selection (present in both modern populations) and divergent selection (present only in one recent breeding population) (Fig. 5A). Around half the sweeps identified in modern populations were early domestication sweeps (97 sweeps, 6.57% of the genome), but with continuously higher selection pressure in the diversification era (Fig. 5B). There were 36 sweeps (2.39% of the genome) selected in parallel in both breeding populations, but not in heirloom varieties, which might relate to novel domestication traits adapting to new farming practices in annualized culture. Equal numbers and sizes of divergent sweeps were found for the two breeding populations (Fig. 5A). A significant positive correlation was found between the total sizes of divergent selective sweeps and the differences in ancestral proportions between the two breeding populations across chromosomes (R^2^ = 0.14, Fig. 5C), indicating the role of divergent selection in altering wild allelic origins.

**Figure 5.**
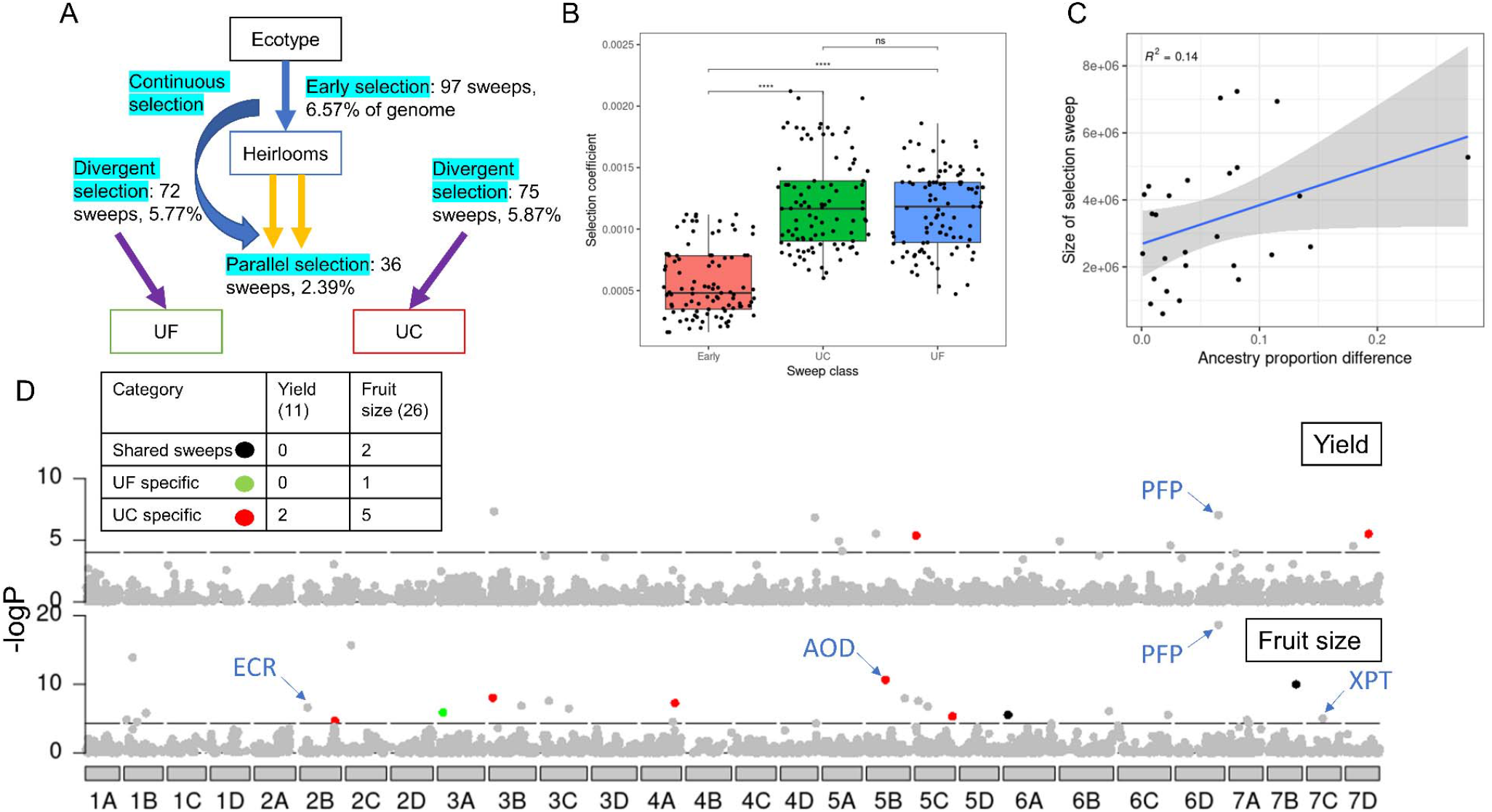
(A) Schematic plot shows the categories of selection sweeps annotated with numbers of sweeps and the total size of sweeps. Three categories include early selection (present in all three scans), parallel selection (present in both populations) and divergent selection (present only in one population). (B) Comparisons of selection coefficient of early domestication sweeps (97 sweeps, 6.57% of the genome) in heirloom and recent breeding populations. The levels of significance are annotated between groups. Four asterisks represent p < 0.00001 (C) Regression of total sizes of divergent selection sweeps against differences in ancestral proportion between two breeding populations across 28 chromosomes. (D) Manhattan plots of genome-wide association study (GWAS) results for yield (gram/plant, upper track, dash line = p_FDR-adjusted_ of 0.05) and fruit size (gram/marketable fruit, lower track, dash line = p_FDR-adjusted_ of 0.01). GWAS signals under selection are colored according to the population it was detected. The breakdown of the number of selected loci is present in the table. The number in parentheses is the total number of GWAS signals for yield and fruit size, respectively. Putative candidate genes are annotated next to the marker positions.

### Genomic loci underling yield (Y) and fruit size (S)

Previously, one fruit firmness QTL^23^ and the *Nerolidol Synthase1* (*FaNES1*) gene^35,36^ were linked to *F. × ananassa* domestication. Here, we expand genotype-to-phenotype associations for two domestication traits using a large-scale genome-wide association study with 1778 individuals. Narrow-sense heritability estimates were 0.434 and 0.698 for Y and S, respectively. Eleven significant signals for Y (p_FDR-adjusted_ < 0.05) and twenty-six for S (p_FDR-adjusted_ < 0.01) (Fig. 5D, Data S2 & S3) were identified. Overlapping GWAS peaks with selective sweeps revealed that only a fraction of the loci was selected by either or both breeding programs (2/11 for Y and 8/26 for S), with a higher number of selected loci in the UCD program (Fig. 5D, Table S6). With the aid of fruit eQTL results and annotations from previous research^36,37^, we identified domestication/diversification -associated genes or genes that could be selected in the future. A strong signal on Chr 6D was co-localized with both Y and S (for Y, p_FDR-ajusted_ = 0.00026; for S, p_FDR-adjusted_ = 1.2E-15), and not under selection with an allele frequency of 0.13 in the GWAS population. A positive marker effect was detected for both traits (β_Y_ = 32.7 gram/plant or more than one fruit per plant, accounting for 2.6% of phenotypic variance; β_S_ = 1.2 gram/fruit, 4.8% of phenotypic variance). Four genes shared cis-eQTL with the GWAS signals (AX-123359105 and AX-123359107), with a pyrophosphate-pyrophosphate-fructose-6-phosphate-phosphotransferase (PFP, FxaC_24g51220) gene having the strongest association (p = 2.93E-11, Fig. 5D, Fig. S14, Table S6). Its paralog in rice regulated carbon metabolism during grain filling, and its rice mutant exhibited reduced grain weight and starch content and increased protein content^38,39^. Additionally, three candidate genes were identified for S. FxaC_18g22380 (lead marker: AX-123361625, Fig. S14, Table S6), a putative acetylornithine deacetylase (AOD) was under selection only in the UCD population with a significant drop of 7 [within the genic region (Fig. S14, Table S6). In *Arabidopsis*, AOD catalyzed ornithine production, and the reduced expression of AOD resulted in early flowering and impaired fruit set^40^. Two other genes/GWAS signals, FxaC_6g07390 (lead marker: AX-123365309, Fig. S14, Table S6), predicted to be a very-long-chain enoyl-CoA reductase (ECR), and FxaC_27g09810 (lead marker: AX-166518420, Fig. S14, Table S6), a likely xylulose 5-phosphate/phosphate translocator (XPT) were not under selection.

### Utilization of extant diversity in modern breeding

Since a huge loss of rare alleles from ecotype to modern populations was observed, introgressing wild alleles not only replenishes genetic diversity, but introduces beneficial traits missing in the current genetic pool. Some successful stories of introgression of wild materials have been reported in strawberry: backcrossing to *F. virginiana* introduced perpetual flowering into UCD varieties, which revolutionized the summer strawberry industry^41^, and introgressing wild genetics improved the nutritional value of a leading European cultivar^42^. However, the speed of recovery for fruit quality traits and retained genetic composition of the wild donor genome in each backcross generation are not known for these examples. Multi-generation modified backcrosses were conducted with *F. virginiana* subsp. *virginiana* (PI 612498) and *F. chiloensis* subsp. pacifica (PI 551736, 48.8% pacifica based on admixture analysis). One representative selection from all generations showed IBD composition close to theoretical estimates from F1 (50%) to BC3 (6.25%) (Fig. S16, Supplementary information). A slightly lower percentage of wild introgression was observed in the *F. chiloensis* backcross (Fig. S16). A positive correlation was found between chromosomal fractions of *F. chiloensis* IBD in BC3 and chromosomal proportions of *F. chiloensis* ancestry in the UF population (Fig. S17, R^2^ = 0.10). Only four generations were required to recover fruit size and sugar content to the level of modern varieties (25 g/fruit, 7.5% for SSC)^43,44^ in those two backcrosses. Specifically, BC3s recovered elite-level average fruit size and sugar content (Fig. S18 & S19) while preserving 6.1% and 7.4% IBD contributions from the wild donors (Fig. S16).

## Discussion

On the basis of genomic analyses of 289 diverse octoploid Fragaria individuals, our ancestral inferences added clarity to the history of strawberry domestication. On a subspecies level, genomic signatures confirm that the initial hybridization occurred between *F. chiloensis* subsp. chiloensis and *F. virginiana* subsp. *virginiana*. Later introgression of *F. virginiana* subsp. *virginiana*, leading to its higher overall genetic composition, and *F. chiloensis* subsp. pacifica happened after migration to NA, confirming past literature and obscure breeding records^1,15,17^. Later introgression of both native NA species likely improved adaptation of new varieties to warm environments^1,17^. In contrast to the pedigree records^15^, there was an absence of *F. virginiana* subsp. platypetala introgression in modern varieties, which implies the introgressed individuals were lost in dead-end pedigrees. Aside from wild hybrids, four octoploid subspecies were distinguished in the phylogenetic and admixture analyses, consistent with previous results using molecular markers^5,6^. Although there were less than 10 individuals of *F. chiloensis* subsp. *lucida, F. virginiana* subsp. grayana, or *F. virginiana* subsp. glauca in the collection, none of them were clustered together, but distributed across another clade.

Like date palm^45^, banana^46^ and citrus^11^, hybrid origin played a major role in the domestication of octoploid strawberry. Genome-wide nucleotide diversity in octoploid strawberry ecotypes (π = 0.0085) was higher than wild populations of other perennial species in the genus Rosaceae such as Prunus (peach, π = 0.0061)^47^, Malus (apple, π = 0.0043)^48^ and even Fragaria nilgerrensis (π1=10.00567)^49^. The burst of diversity and effective population size within a few founder individuals due to interspecific hybridization provided immense phenotypic variation for recurrent selection. Following the initial hybridization, strong selection pressure stimulated a rapid decline of Ne lasting around ten generations, leading to a severe bottleneck (Fig. 4B). This Ne inference is in line with pedigree-based sociogram analysis for the UCD population, as a narrow channel occurred between 1940 and 1980^15^. The UF population was heavily rooted in UCD varieties released between 1950 and 1980 (Fig. S10), and almost identical patterns were revealed in the demographic trajectory for both populations. The estimated range of total generations of breeding (10 to 20 generations) was consistent with previous estimations using genealogy data (16 years/generation)^15^.

The two breeding populations preserved 0.36 and 0.26 of total alleles from ecotypes but a lower 0.19 and 0.09 of rare alleles (MAF<0.2). Even in the heirloom varieties, 71 percent of rare alleles were lost due to the small number of founders. These estimates should be reliable due to a large number of wild individuals (n=105) and the non-mapping-required kmer-based diversity inference approach that captured alleles within structural variants, which are pervasive among the octoploids^36,50^. The greater diversity of the UF population compared to UCD also seems reasonable, during the studied timeframe, due to known admixture in the UF population from multiple resources, such as southeastern U.S. germplasm, UCD varieties developed 1950 to 1980^51^ and Australian germplasm^52^. These various resources were utilized to combine resistance to fungal pathogens and other subtropical climatic adaptations with improved yield and quality. During this regional adaptation, chromosomal proportions of two ancestral species were differentially selected for two breeding programs. ^9,10^The selection for different ancestral proportions in multiple chromosomes might be linked to adaptive traits such as heat tolerance, flowering time, and chilling requirement, which warrants future QTL studies.

In strawberry, diversity can be rapidly recovered through introgression from wild relatives^8^. In backcrosses with both wild progenitor species, BC3 individuals recovered elite fruit size and soluble solids, while still preserving around 6.25% of wild alleles IBD. An alternative approach has been sought to boost genetic diversity via hybridization of individuals with superior horticultural traits from the two wild octoploid species^53^. Compared to the reconstruction approach, backcrossing is more rapid and targeted, and requires a smaller number of progenies to screen, as genetics of elite germplasm is mostly preserved, but limits the amount of wild diversity that can be incorporated in each designed backcross.

Diversification and domestication are always accompanied by multiple phenotypic changes. One of the reasons that seed crops often experience protracted domestication history is that domestication traits such as non-shattering and seed size often underwent unconscious selection with low levels of selection pressure in early domestication^54^. Unlike seed crops, aside from dramatic quantitative changes in plant vigor, yield, fruit size, and fruit firmness, the cultivated strawberry and its wild progenitors do not differ in any obvious qualitative way^4^. This allowed a strong and conscious selection on fruit size, firmness, and yield, leading to a rapid domestication process. Using the large GWAS population, eleven and twenty-six loci were found associated with yield and fruit size, respectively. More than half of the loci were not under selection, including a locus with a strong directional effect on both traits (Fig. 5D). Discovery of its putative causal gene pyrophosphate-pyrophosphate-fructose-6-phosphate-phosphotransferase (PFP, FxaC_24g51220) warrants future investigation into its function in fruit production.

Strawberry represents an ideal model to study crop domestication, as it encapsulated hybridization, introgression and divergence in a short domestication history. Hybridization and introgression supplied incredible amounts of genetic diversity and phenotypic variation within interfertile species. Thereafter, strawberry breeders rapidly improved fruit quality and yield within twenty generations by recurrent selection with a few targeted introgression from octoploid progenitor species. Species-preferential selection of the improved hybrid rendered varieties adapted to annualized culture in Mediterranean and subtropical climates. While there is substantial overall diversity in cultivated strawberry, rare alleles have been largely purged, suggesting a targeted introgression approach for specific, desired traits may be the best breeding approach for the use of octoploid wild species in the future. Meanwhile, the loci identified here underlying fruit size and yield should be explored further to facilitate additional improvements in these traits.

## Materials and methods

### Sequencing and variant calling

Illumina WGS was obtained for 290 octoploid strawberry individuals, doubling the number from the previous WGS study^23^, including 68 accessions from the University of California, Davis (UCD), developed between 1988 and 2011^23^; 52 accessions from the University of Florida (UF) developed between 2009 and 2019 (besides ‘Florida Elyana’); 65 heirloom varieties including 32 varieties developed outside of the US or other US programs released before 1990, 17 UCD accessions developed before 1990, nine UF accessions developed before 2000, and seven F_1_ and BC_1_ wild octoploid introgression lines developed at UF; 43 Fragaria chiloensis accessions; 59 Fragaria *virginiana* accessions; and three Fragaria × ananassa ssp. cuneifolia accessions (Table S1). Among them, 80 individuals, including most founders and modern varieties of the UF breeding program, were newly sequenced with an average output of 30.5 Gb, equivalent to ∼39× of the octoploid strawberry genome. The rest were retrieved from NCBI (Bioproject Accessions PRJNA402067, PRJNA57838 and PRJNA727900) published from three previous studies^23,26,55^. The entire dataset contained 5.16 Tb, with 17.7 Gb per sample, equivalent to ∼23×. Genomic DNA was isolated from young leaves using a customized CTAB method. After initial QC, a 150 bp pair-end (150PE) library was prepared for individual sample. Sequencing was conducted on NovaSeq platform. Service was provided by the UF Interdisciplinary Center for Biotechnology Research, Gainesville, FL, USA. Illumina short reads were aligned to the FaRR1 octoploid genome^21^ using the SNAP aligner v2.0.0^56^ with the “-so” flag to remove duplicated reads and sort bam files. SNPs and InDels were identified by the HaplotypeCaller module in the GATK v4.1.9^57^ with the gVCF mode following best practices. Then GenomicsDBImport module merged GVCFs of all samples before jointly genotyping via the GenotypeGVCFs module with the “--genomicsdb-shared-posixfs-optimizations” flag. Variants were initially hard filtered using VariantFiltration module with the parameters “QUAL < 30; QD < 2; FS > 55; SQR >3; MQ < 55; MQRankSum < -2”. Additional filtering of missing fraction < 0.3 was conducted using bcftools v1.15. SNPs and InDels failing QC were removed. The final variant set included 94,944,176 SNPs and 15,316,300 InDels. Variants were annotated using annovar v20191024^58^. One individual, CFRA_1927, with low coverage was removed for further analysis. A SNP dataset (MAF > 0.01) was extracted from the full dataset, containing 23,833,040 SNPs.

### PCA, *Fst*, admixture and demographic analyses for wild octoploid species

The SNP set was further pruned using R^2^ < 0.4 in non-overlapping windows of 1000 SNPs and intronic regions. A total of 259,491 SNPs were retained for PCA and admixture analysis. Principal component analysis was performed using SNPRelate v1.26 in R^59^. Admixture analysis was conducted using fastSTRUCTURE v20150112 with the default parameters^60^. Between 3 and 10 K were tested, with K = 7 maximizing marginal likelihood. K = 3 was also plotted due to clear separation of three species. A neighbor-joining (NJ) tree for all individuals was constructed using VCF-kit v0.2.6. Effective population size trajectory and split time estimation were computed among *F. chiloensis* and F. *virginiana, F. virginiana* subsp. *virginiana* and *F. virginiana* subsp. platypetala, *F. chiloensis* f. patagonica and *F. chiloensis* subsp. pacifica using SMC++ v1.15.4^61^ with the full SNP dataset. A mutation rate of 2.8 × 10^−9^ per site per year was based on the previous estimation of Fragaria species^62^. The confidence interval of the split time was computed as 5% and 95% quantiles across estimations for individual chromosomes. More complex demographic models were built on folded allelic frequency spectrum (AFS) via Dadi v2.1.1^30^ using 1,847,930 biallelic SNPs that were randomly selected from every ten consecutive SNPs from the full SNP set. Admixture individuals were removed for the demographic inference, and 42 and 38 *F. chiloensis* and *F. virginiana* were retained for the analyses. Three Dadi models were tested: species formation with asymmetric migration; species formation with exponential growth and asymmetric migration; and species formation followed by an initial stage of equilibrium with asymmetric migration, and then exponential growth and asymmetric migration. Every model was simulated 100 times. The model with the highest likelihood was selected and residual AFS plots were visually examined.

### Species tree and introgression test

A maximum likelihood (ML) species tree was built including seven diploid Fragaria species with multiple individuals within F. vesca subsp. vesca (n = 3) and F. vesca subsp. bracteate (n = 2)^20^, four octoploid species showing separation in the admixture analysis with four individuals in each, and Potentilla anserina as an outgroup (Table S2). The Illumina short read data of diploids was retrieved from multiple studies ^20,62–64^. All the wild octoploid individuals were free of admixture according to the admixture analysis. To overcome the problem of mapping species with different ploidy levels, diploid samples were mapped to subgenome A of the FaRR1 genome^21^, while octoploid samples were aligned to the entire genome. The variant calling and filtering for only subgenome A was identical to the aforementioned. A total of 1,866,574 biallelic SNPs on subgenome A was concatenated for phylogeny inference. A ML tree was built using the GTR+F+ASC+R3 model in IQ-TREE v2.2.2 with the parameters “-alrt 1000 -B 1000” for 1000 replicates of SH-like approximate likelihood ratio test and ultrafast bootstrap approximation^65^. Introgression tests including D, f4-ratio, f-branch statistics among all Fragaria species were analyzed in Dsuite v0.4^66^.

### Ancestral proportion estimates and demographic inference for *Fragaria × ananassa*

Local ancestry for all *F. × ananassa* varieties was inferred using RFMix v2.03^31^. Samples of *F. chiloensis* f. chiloensis and *F. chiloensis* f. patagonica, *F. chiloensis* subsp. pacifica and *F. chiloensis* subsp. *lucida* samples, and *F. virginiana* subsp. *virginiana* and *F. virginiana* subsp. grayana were separately grouped and regarded as ancestral populations. Two rounds of phasing were conducted, first with Beagle v5.2^67^ and then SHAPEIT v2.12^68^. The phased vcf file was provided to RFMix. To infer the trajectories of effective population size (Ne) for breeding populations over the past three hundred years since the initial hybridization, a linkage disequilibrium-based demographic history inference approach was implemented in the software GONE^69^. The analysis was carried out separately for the UF (n = 52) and UCD accessions (n = 68). The full SNP set was used as input and recombination rate was approximated at 0.25 Mb per cM. ALDER v1.03^32^ was also used to estimate the time of hybridization with the two references mode, provided with SNP data of both *F. virginiana* and *F. chiloensis*.

### Identical by descent (IBD), inbreeding coefficient, runs of homozygosity, nucleotide diversity, and K-mer-based diversity estimates

IBD percentages of all octoploid individuals were computed using the Plink method implemented in the SNPRelate v1.26 package in R with the LD-pruned SNP set (N = 259,491). Inbreeding coefficient (F) was measured using vcftools v0.1.16 with the flag “--het” and the full SNP set. Long runs of homozygosity (LROH) for each individual were measured using “roh” mode in bcftools v1.15 with the full SNP set. Nucleotide diversity (π) was calculated using vcftools v0.1.16 in windows of 100 kb for each group with the full marker set. Ecotype was a joint set including both *F. virginiana* and *F. chiloensis* individuals.

To investigate diversity changes during domestication and diversification, a customized k-mer pipeline was built, modified from a previous study^70^. K-mers (k = 31) with a minimum count of three were first counted in raw short reads data for each individual using KMC v3.2.1^71^. The distribution of total k-mer counts derived from raw read was benchmarked with k-mer counts of published phased genomes^21,36^. The distribution of sample k-mer counts was centered between the phased assemblies of FaRR1^21^ and the FL 15.89-25^36^, demonstrating direct k-mer counts from raw data were as adequate as counts from assembled genomes. Next, entries were recoded to 1 (presence) and 0 (absence). K-mer counts of samples in each group were summed. Within each group, k-mers occurring in less than two individuals were removed. The core k-mer set for *F. × ananassa* was estimated using 15 samples with the highest sequencing coverage in UF, UCD and heirloom sets, and core k-mers were removed from each group. The intersecting k-mer sets between each pair of groups were computed. Retained percentage of diversity was computed as 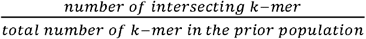.

### Identification of selective sweeps

Selective sweeps were identified using XP-CLR^72^ in windows of 5000 bp. Early selective sweeps were detected for the ecotype (n = 92) to heirloom (n = 40) transition. Only heirloom varieties developed before 1980 were included to avoid detection of sweeps in the diversification era. The selective sweeps for two breeding populations adapted for Mediterranean and subtropical climates were identified comparing ecotype to UCD (n = 68) or UF (n = 52) varieties and breeding selections, which were an ensemble of early domestication sweeps, parallel sweeps and divergent sweeps. The top 5% of windows with the highest selection coefficients were selected, and signals within 400 kb were merged based on the previous estimation of LD decay in modern varieties^23^. The highest selection coefficient within a selective sweep was used to represent the selection pressure of that sweep. A customized R-script was used to compile XP-CLR output and identify selective sweeps. Only the total size of divergent sweeps was correlated with the difference in chromosomal ancestral proportion between the two programs.

### Genome-wide association study (GWAS) for marketable yield and fruit size

A total of 1778 individuals representing the elite genetic pool of the UF breeding program were trialed over five consecutive seasons (2016-2021). Data from seasons 2016-17 and 2017-18 were previously used for validation of genomic selection models^73^. Replicated trials were established at the Gulf Coast Research and Education Center in Balm, FL (lat. 27° 45′ 37.98″ N, long. 82° 13′ 32.49″ W). Five replicates with single plant per replicate were arranged in a randomized complete block design. The bare-root plants were planted in raised beds in the early October and managed according to standard commercial practices. Every year, between 411 (2016-17 season) and 452 (2017-18 season) genotypes were planted, and between 67 to 140 common genotypes were replicated across consecutive years. Yield (Y, gram/plant) and fruit size (S, gram/marketable fruit) were measured for 16 weeks from the third week of November to the first week of March. Best linear unbiased estimates adjusted for year effects (Model: Y/S = Genotype + Year + Error) were the input for GWAS models. All individuals were genotyped with either iStraw 35k^74^ or FanaSNP 50k SNP arrays^22^; the common 5895 markers between two arrays were used for GWAS. The marker positions were anchored to the FaRR1 genome based on blast results. The GWAS method, FarmCPU^75^ implemented in GAPIT3 R package^76^ was used for both traits with the inclusion of the top five principal components and markers with minor allele frequency > 0.05. Phenotypic variance explained by a marker was computed as the difference of R^2^ in general linear mixed models with and without the target marker. To overlap with selective sweeps, a GWAS locus was defined as a window of 400kb flanking both sides of the lead marker. The putative causal genes underling GWAS signals were determined based on genes sharing the same cis expression QTL in a previously developed fruit eQTL map^36^. Only GWAS signals with fewer than seven linked genes were further investigated based on gene annotations.

### Backcrosses and field measurements

Two backcross families for *F. virginiana* subsp. *virginiana* (PI 612498) or *F. chiloensis* subsp. pacifica (PI 551736, partially admixed) were initiated in 2011 and 2012, respectively. Selection was conducted in every cross and season based on visual observations of morphological characteristics. In every week from 12/07/21 to 1/18/22, average fruit weight was measured for one representative selection of the wild parent, and F1, BC1 and BC3 generations for the *F. chiloensis* backcross, as well as the wild parent, and BC1, BC2 and BC3 generations of the *F. virginiana* backcross. All fruits from ten clones in a single plot at the Gulf Coast Research and Education Center for each individual were collected. Brix% (soluble solids content) of fruit was measured using a handheld digital refractometer in weeks two, six and seven. The mean reading from two fruits for each individual was recorded. A customized kmer-based pipeline was built to infer IBD segments. Details of the pipeline can be found in the supplementary information.

## Supplementary table legends

Table S1. Sample information of 290 re-sequenced individuals.

Table S2. Individuals that are used in max-likelihood species tree.

Table S3. Number of variants in each category.

Table S4. Estimation of parameters in the best-fit dadi model

Table S5. Proportion of Fragaria *virginiana* ancestry across different breeding populations

Table S6. GWAS signals for two traits and candidate genes associated with the markers. The following sheets detail all the associated genes to the marker based on previous eQTL mapping results.

## Acknowledgements

We thank Dr. Steven Knapp and Dr. Mitchell Feldmann from the University of California, Davis for reviewing early versions of the manuscript. We thank Dr. Doug Soltis and Dr. Pam Soltis from the University of Florida for their advice on the phylogenetic work and review of the manuscript. We thank Dr. Nahla Bassil from USDA for providing pictures of wild species. We thank the strawberry breeding programs at UCD and UF for their assistance in sample collection. We thank UF research computing for computational resources and technical assistance and the Florida Strawberry Research and Education Foundation for their generous funding.

## Authors’ contributions

ZF and VMW conceptualized the study. ZF conducted analyses. ZF and VMW wrote the manuscript.

## Data Availability

Raw whole genome sequencing data for 80 newly sequenced individuals is deposited in the NCBI under accession number PRJNA986921. SNP map in VCF.gz format and raw data for GWAS study is available at Zenodo with DOI: 10.5281/zenodo.8067127.

