## Supplementary Figures for "Genomic signatures of strawberry domestication and breeding"

### Supplementary information

**Identical-by-descent (IBD) segment inferences**

The pipeline started by counting 31-mers for all samples and subtracting the aforementioned core set of kmers representing the modern strawberry genetic pool from each individual using KMC v3.2.1[47]. Next, the IBD kmers were extracted from each progeny by intersecting its kmer set with the corresponding wild doner's kmer set. After that, the FASTQ file of each progeny was filtered with its IBD kmer set to remove reads that do not contain IBD kmers, and filtered reads were mapped to the FaRR1 genome with the SNAP aligner v2.0.0[29] with the "-so" flag. The read coverage was measured using the count module in the igvtools v2.8.6 with a widow size of 5000bp, and then converted to the bedgraph format. IBD segments in each progeny were identified using a novel algorithm that enabled the auto-detection of the coverage cutoff after spline-smoothing the coverage signals. First, coverage signals were smoothed with the spline smoothing function in the npreg package v1.0 in R. Second, assuming the coverage followed a bimodal distribution, we detected the junction of two peaks by determining the coverage value with the lowest density after removing values beyond 10 and 90 percent quantiles. Third, standard deviation (SD) and mean of the low-coverage (non-IBD) peak were determined and the cutoff was estimated as the mean plus three times SD if the identified junction was beyond or equal to four times SD, or the coverage value at the junction minus one SD. Fourth, IBD segments were identified in windows of 200kb with a step of 100kb. Fifth, given IBD not present in parents should not exist in its progenies, we corrected progenies' IBD based on their parents' IBD. This algorithm was tested for backcross generations from F1 to BC3. Some parameters need to be adjusted when applied to the BC4 generation or beyond, such as the cutoff quantiles. The developed R functions and data were deposited in github.

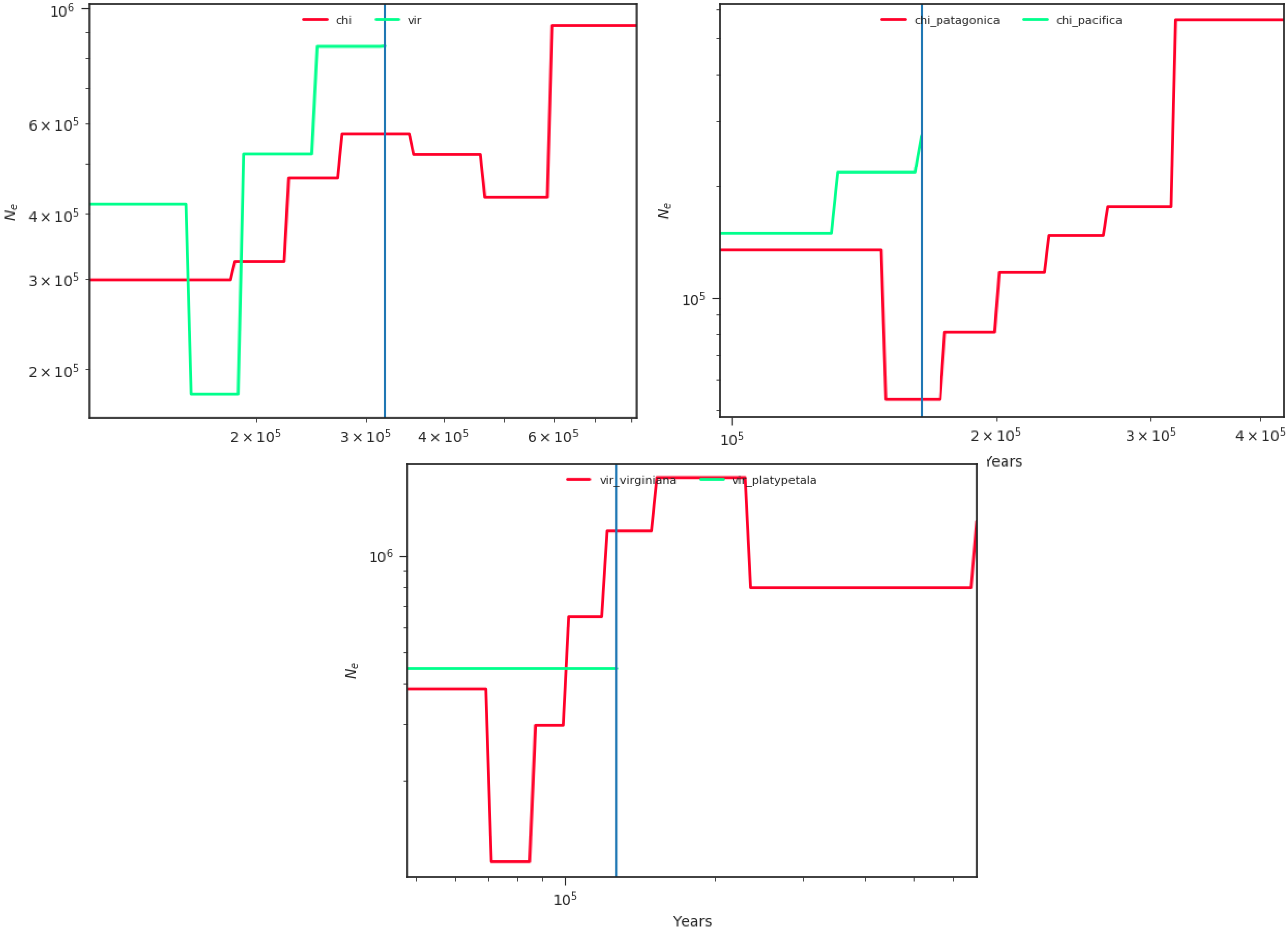


Figure S1. Splitting time and effective population size change inference of *Fragaria chiloenesis* (chi) and *Fragaria virginiana* (vir) and subspecies within either species, computed by SMC++.

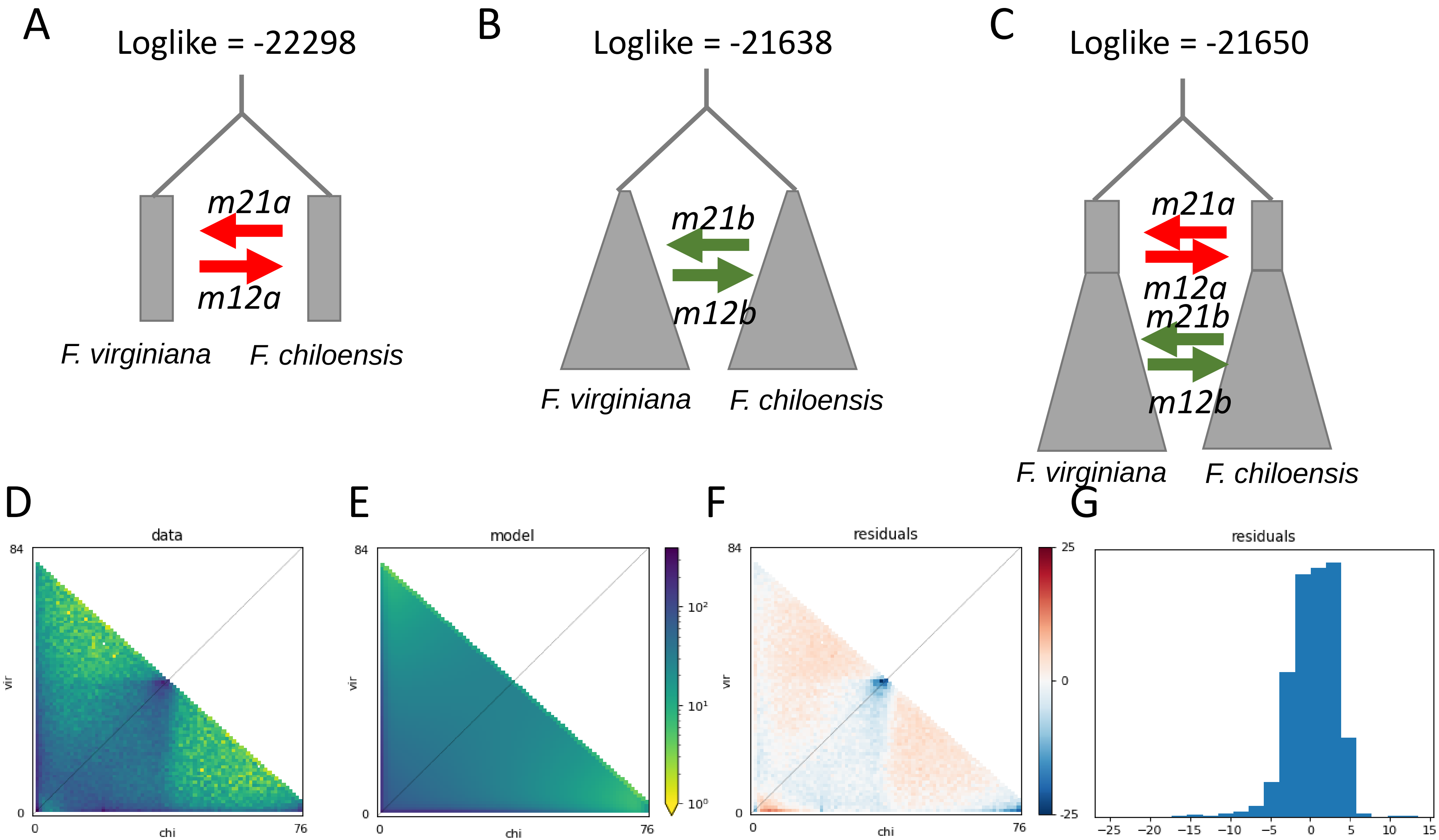


Figure S2. Three dadi models were tested and illustrated. Log likelihood of the models is shown. The model schemes are (A) species formation with asymmetric migration; (B) species formation with exponential growth and asymmetric migration; (C) species formation followed by an initial stage of equilibrium with asymmetric migration, and then a split with exponential growth and asymmetric migration. (D) Sample two-dimensional allelic frequency spectrum (AFS). (E) Simulated AFS, (F) Residual AFS, and (G) Residual distribution based on model (B).

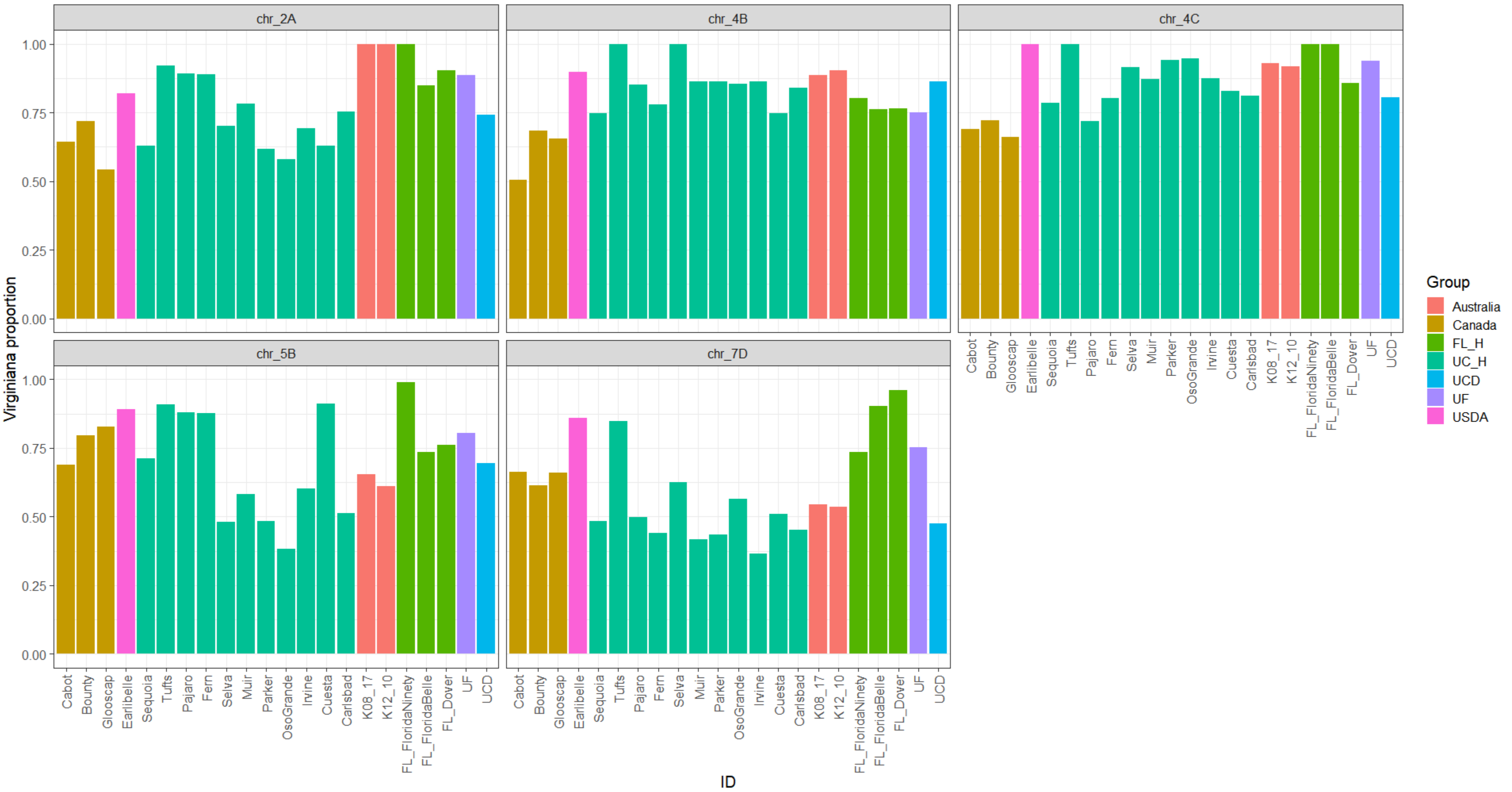


Figure S3. Proportions of Fragaria virginiana ancestry in founders of the University of Florida breeding population. Each panel represents one chromosome. Bars are colored according to the regions or programs of development. FL_H and UC_H represent heirloom varieties from University of Florida and University of California, Davis. Both are in the pedigree for UF varieties.

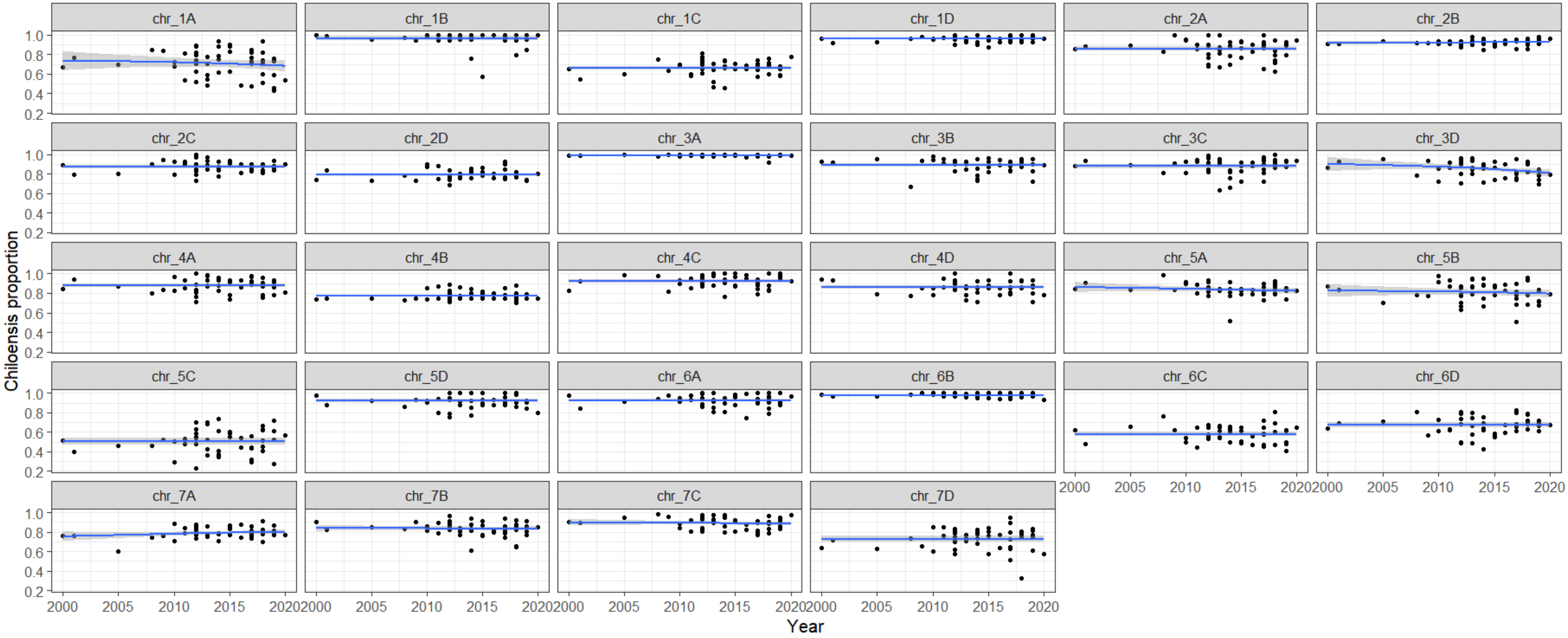


Figure S4. Proportions of *Fragaria virginiana* ancestry across individuals developed between 2000 to 2020 from the University of Florida breeding program. The x axis presents the year of crossing. The best-fit line is plot with confidence interval.

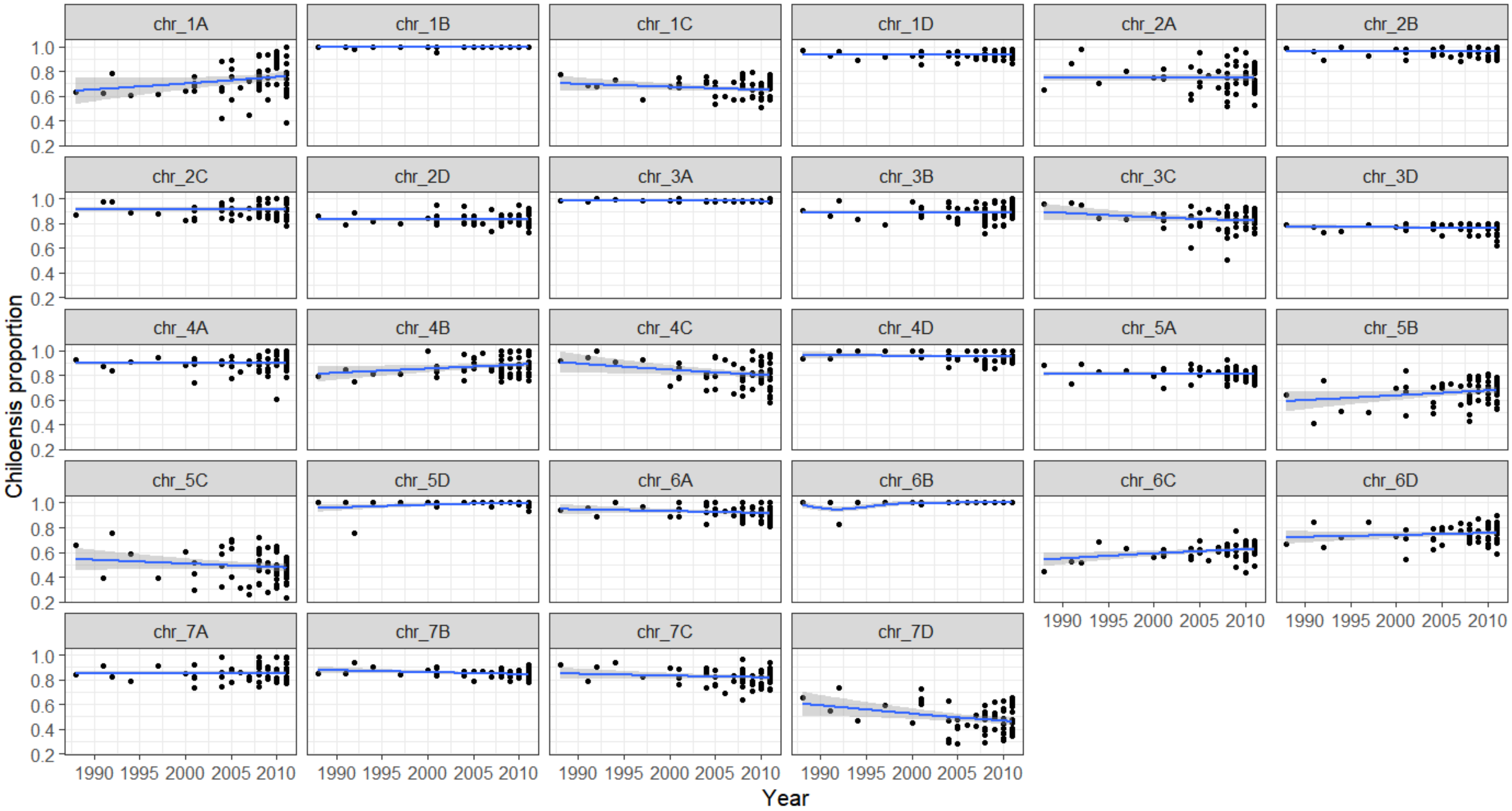


Figure S5. Proportions of Fragaria virginiana ancestry across individuals developed between 1980 to 2010 from the University of California, Davis breeding program. The x axis presents the year of crossing. The best-fit line is plot with confidence interval.

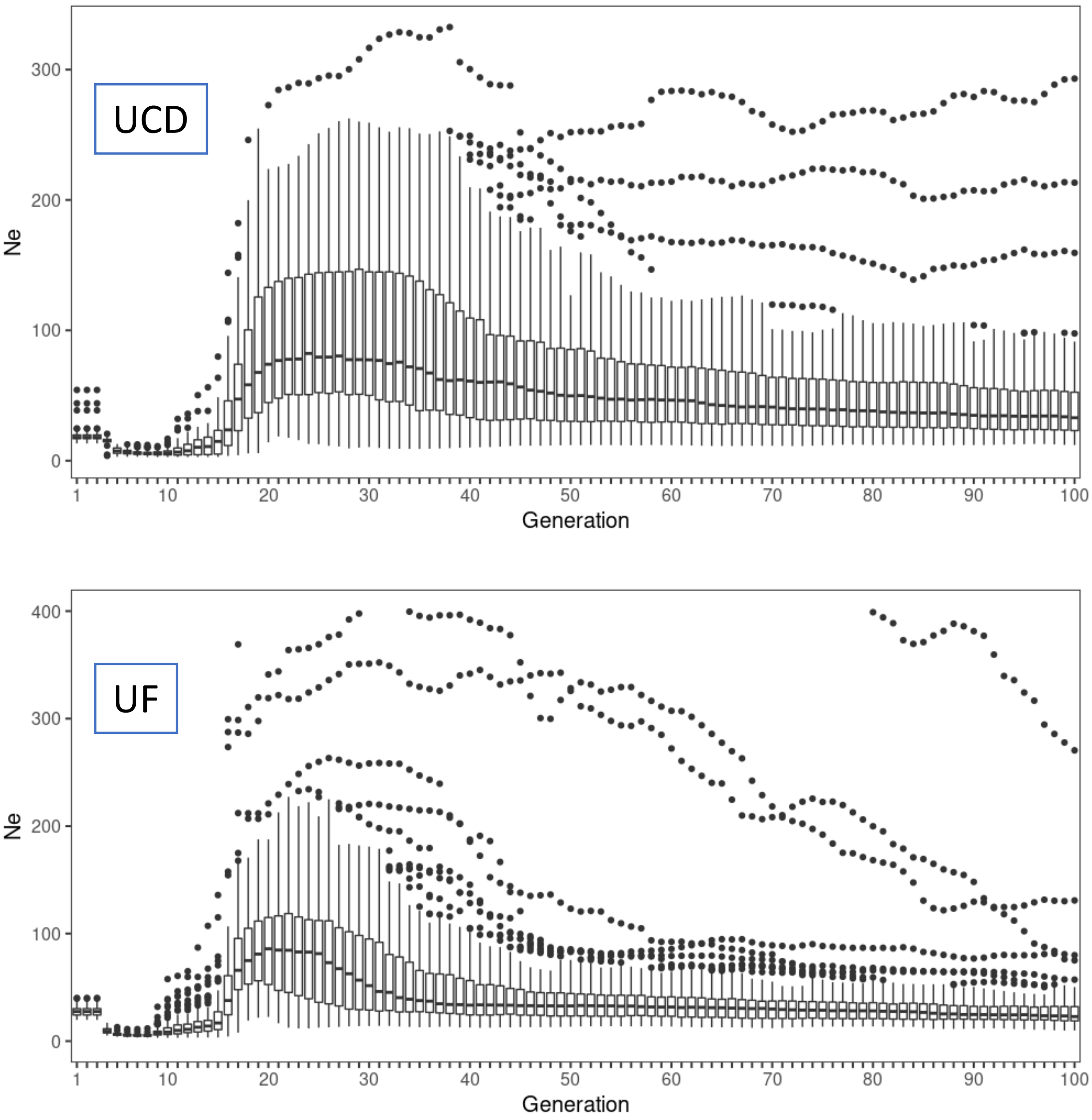


Figure S6, Effective population size (Ne) estimates in last 100 generations for University of California-Davis (top) and University of Florida (bottom) breeding populations. Boxes represent 25 and 75 quartiles of 100 replicates. Middle lines represent medium values.

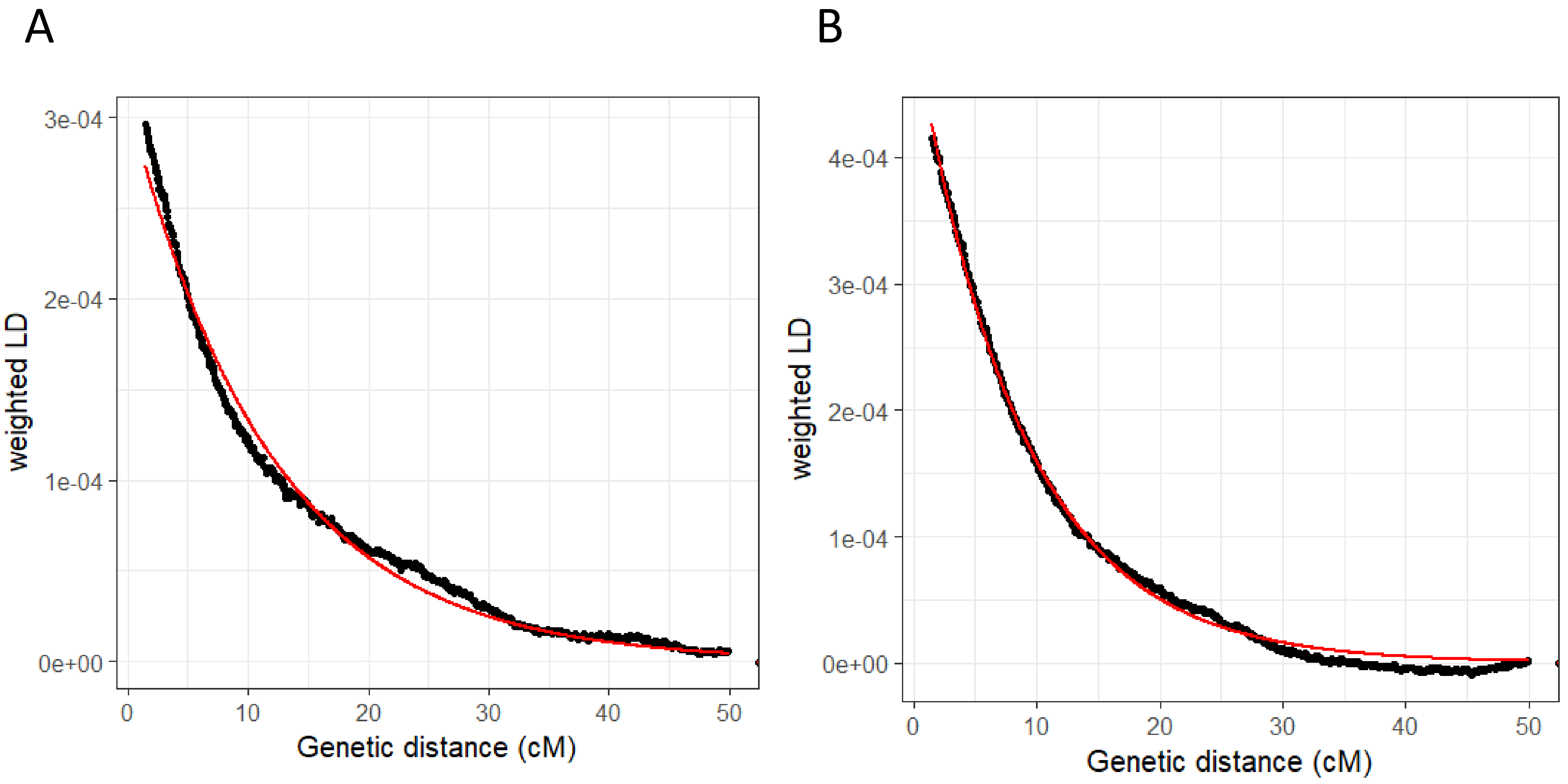


Figure S7. Weight LD decay curves of the University of California-Davis population (A) and the University of Florida population (B). The estimates of Weight LD decay using best-fit exponential decay curve (red) starting from 1.3 cM, are plotted with real data (black).

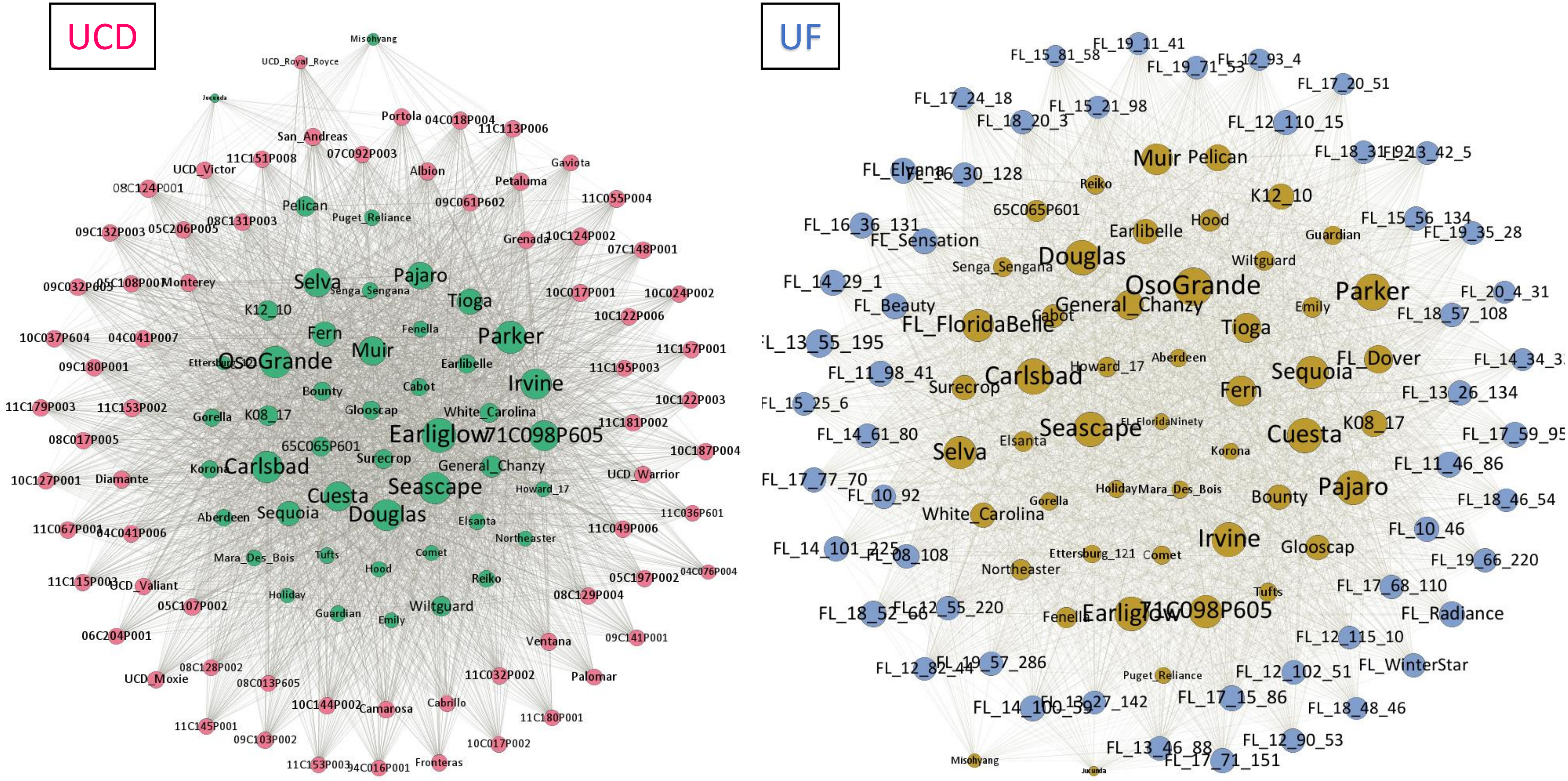


Figure S8. Identical by descent (IBD) networks show the heirloom varieties shared IBD with either University of California-Davis or University of Florida accessions. Only heirloom varieties and accessions from one breeding program are included in corresponding IBD network. The size of dots is proportional to weighted IBD, calculated as the sum of all IBD relationships.

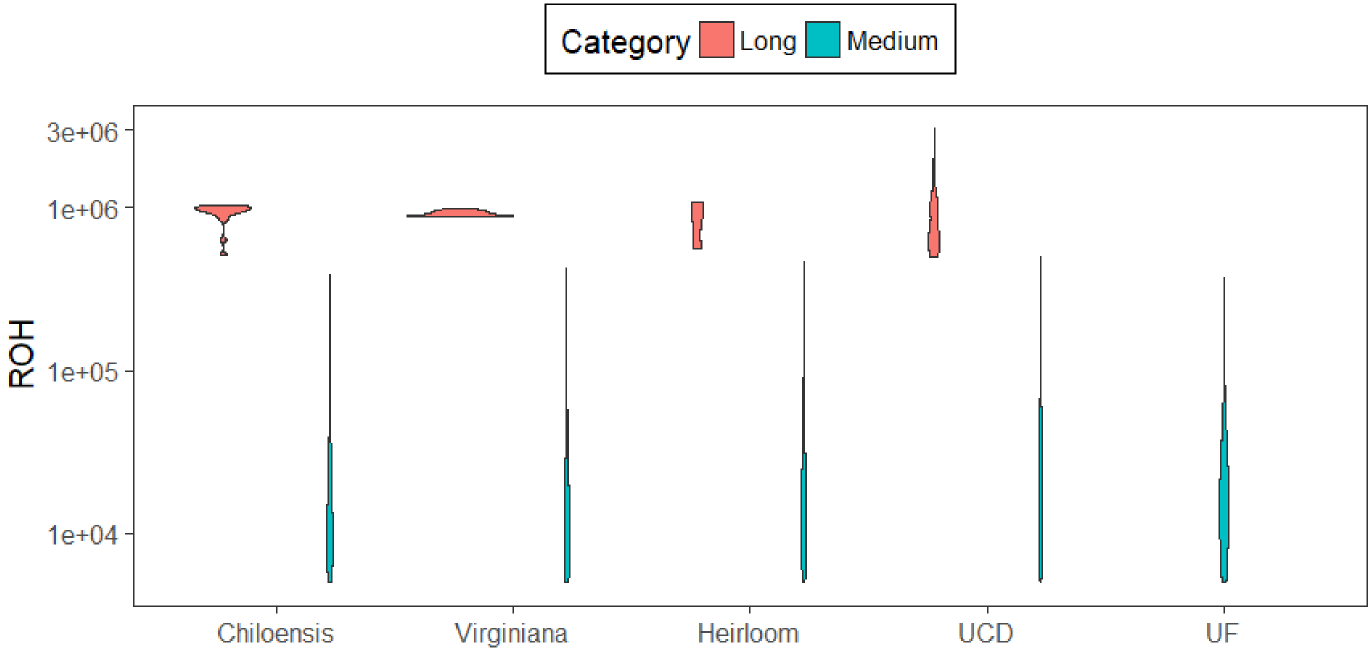


Figure S9. Distributions of runs of homozygosity in different populations. Length distribution is separated into two categories, long (length > 500kb) and medium (500kb > length > 5kb).

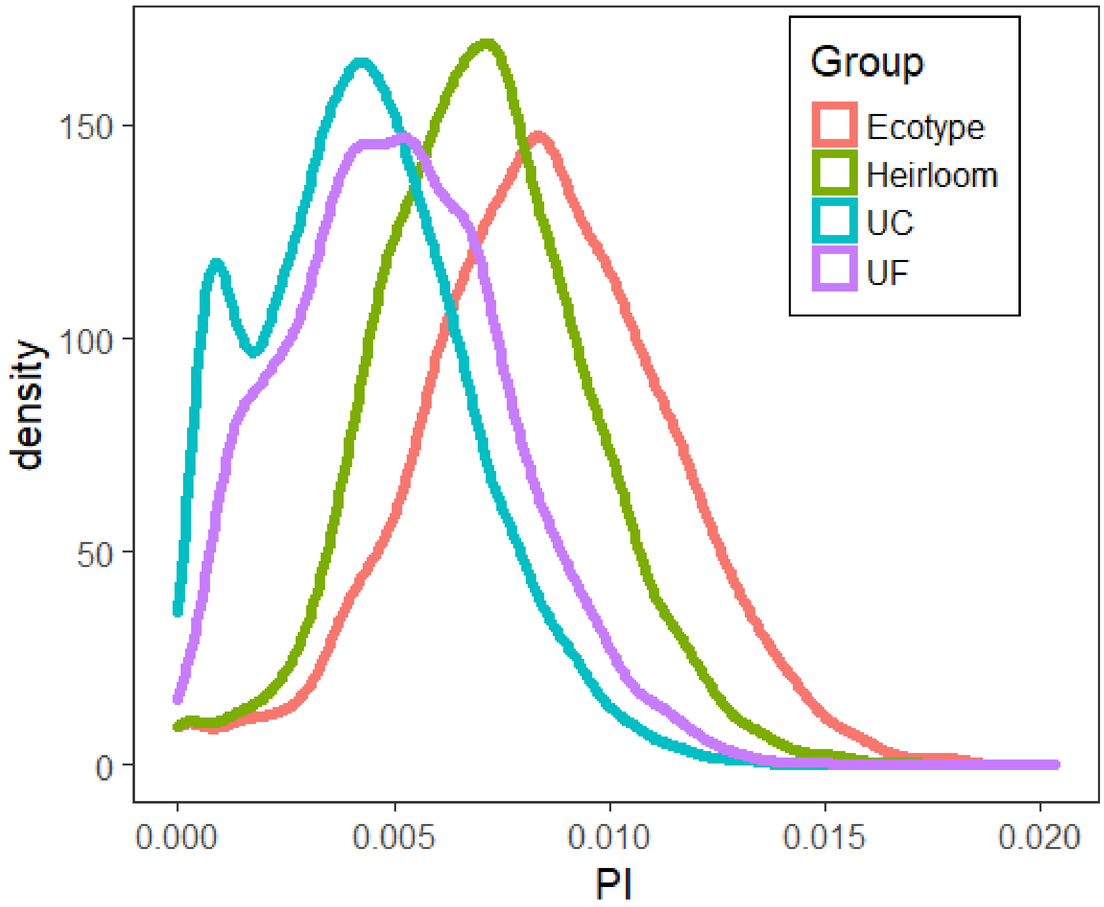


Figure S10. Nucleotide diversity (π) distributions in windows of 100kb across four populations.

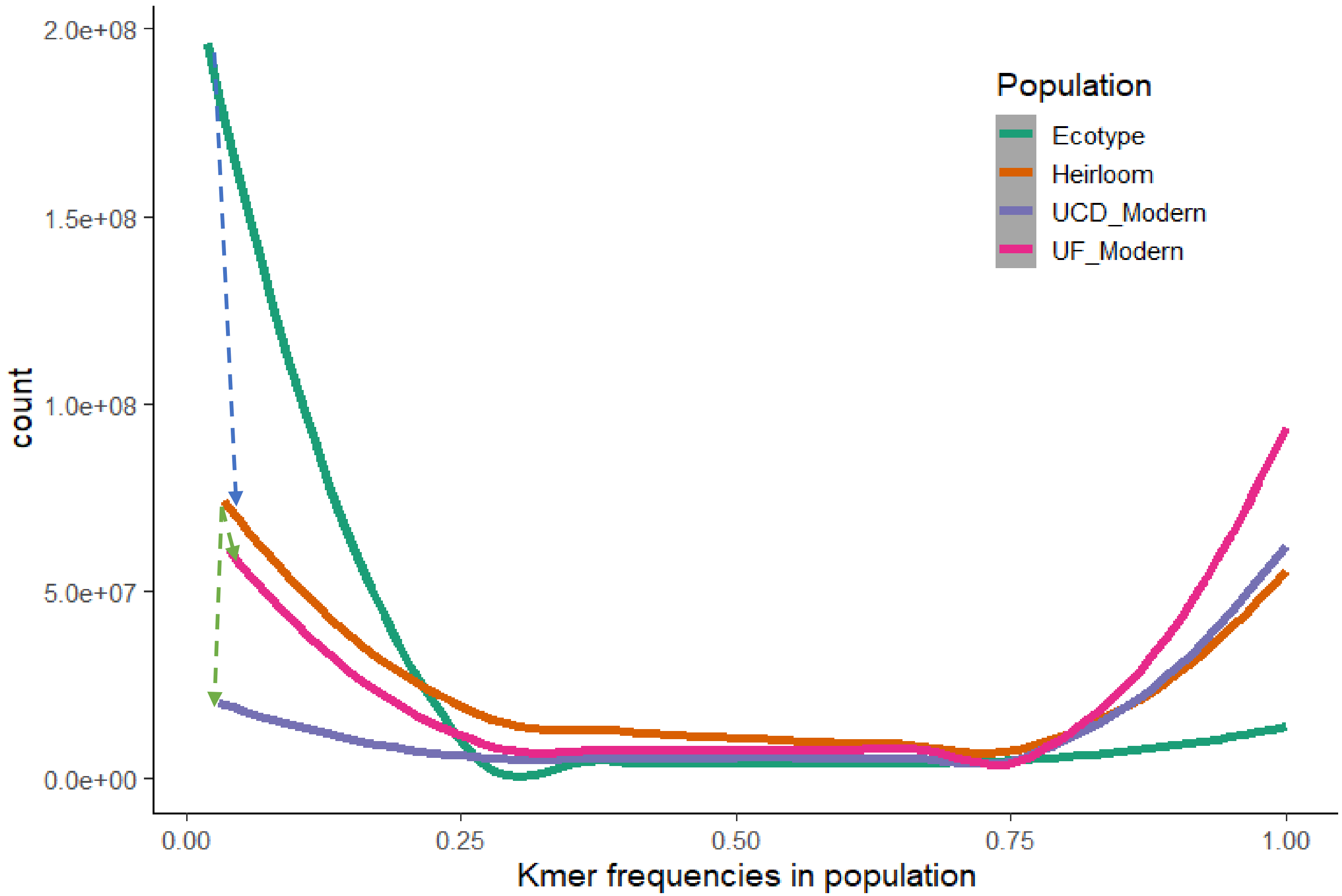


Figure S11. K-mer (k = 31) counts frequency distributions for different populations. Arrows illustrate loss of rare alleles from ecotype to heirloom (blue) and from heirloom to modern accessions (green).

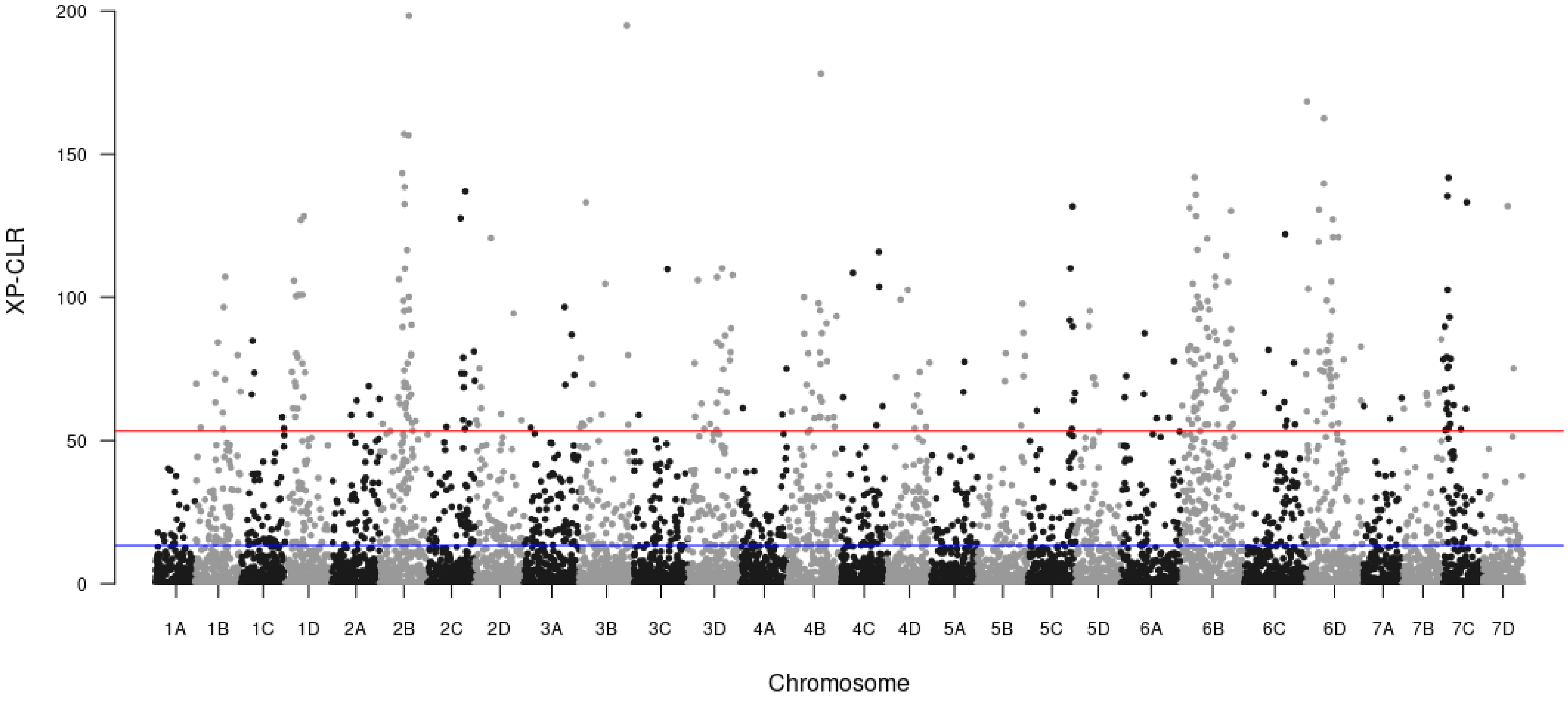


Figure S12. Cross-population composite likelihood ratio (XP-CLR) statistics of sweeps during early domestication using ecotypes and heirloom varieties. The red and blue line represent the upper 0.01 and 0.05 quantiles of XP-CLR estimates.

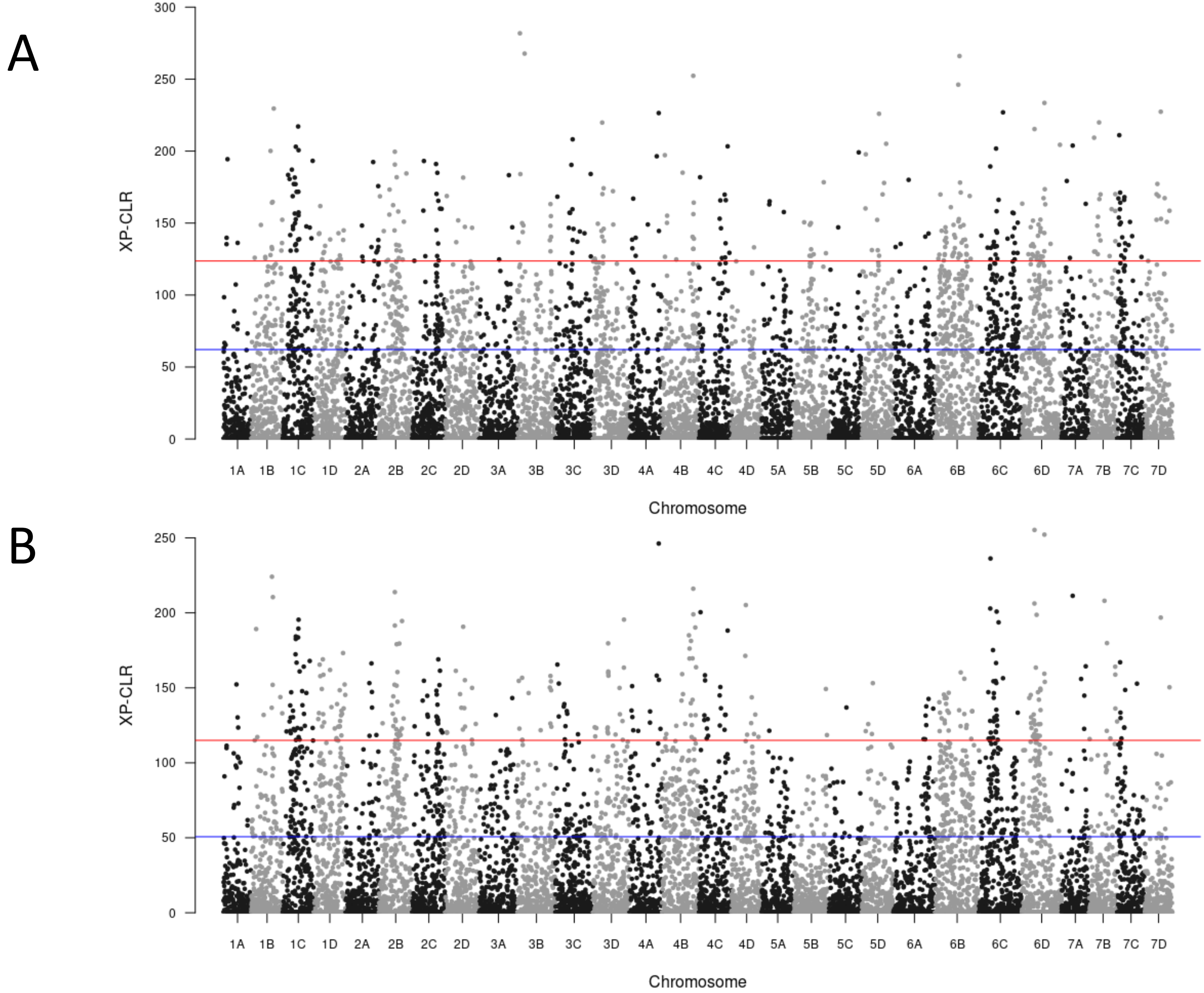


Figure S13. Cross-population composite likelihood ratio (XP-CLR) statistics of sweeps for University of California-Davis (A) and University of Florida (B) populations. The red and blue line represent the upper 0.01 and 0.05 quantiles of XP-CLR estimates.

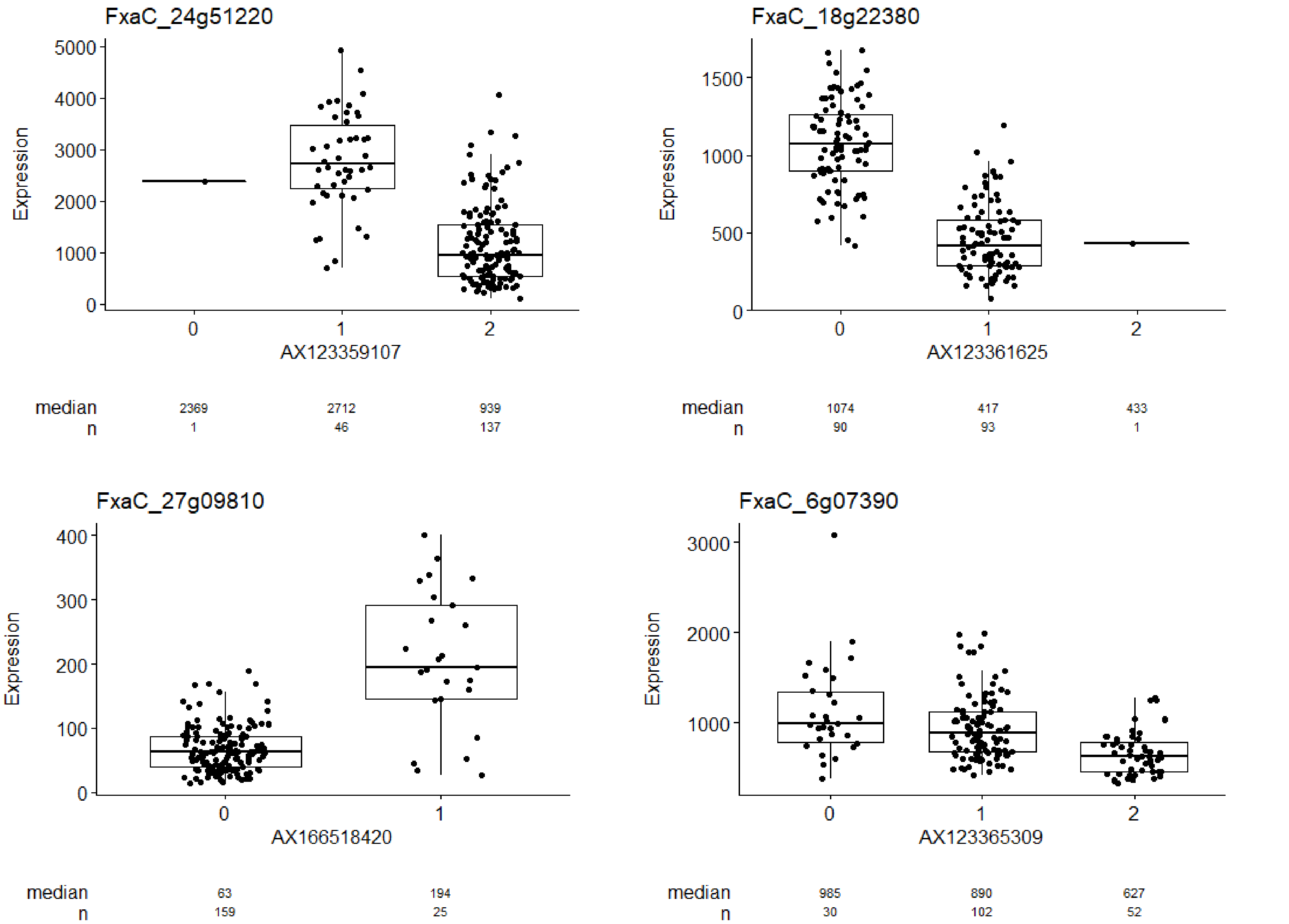


Figure S14. Expression changes for the putative causal genes among different allelic dosages. A dot represents expression for an individual.

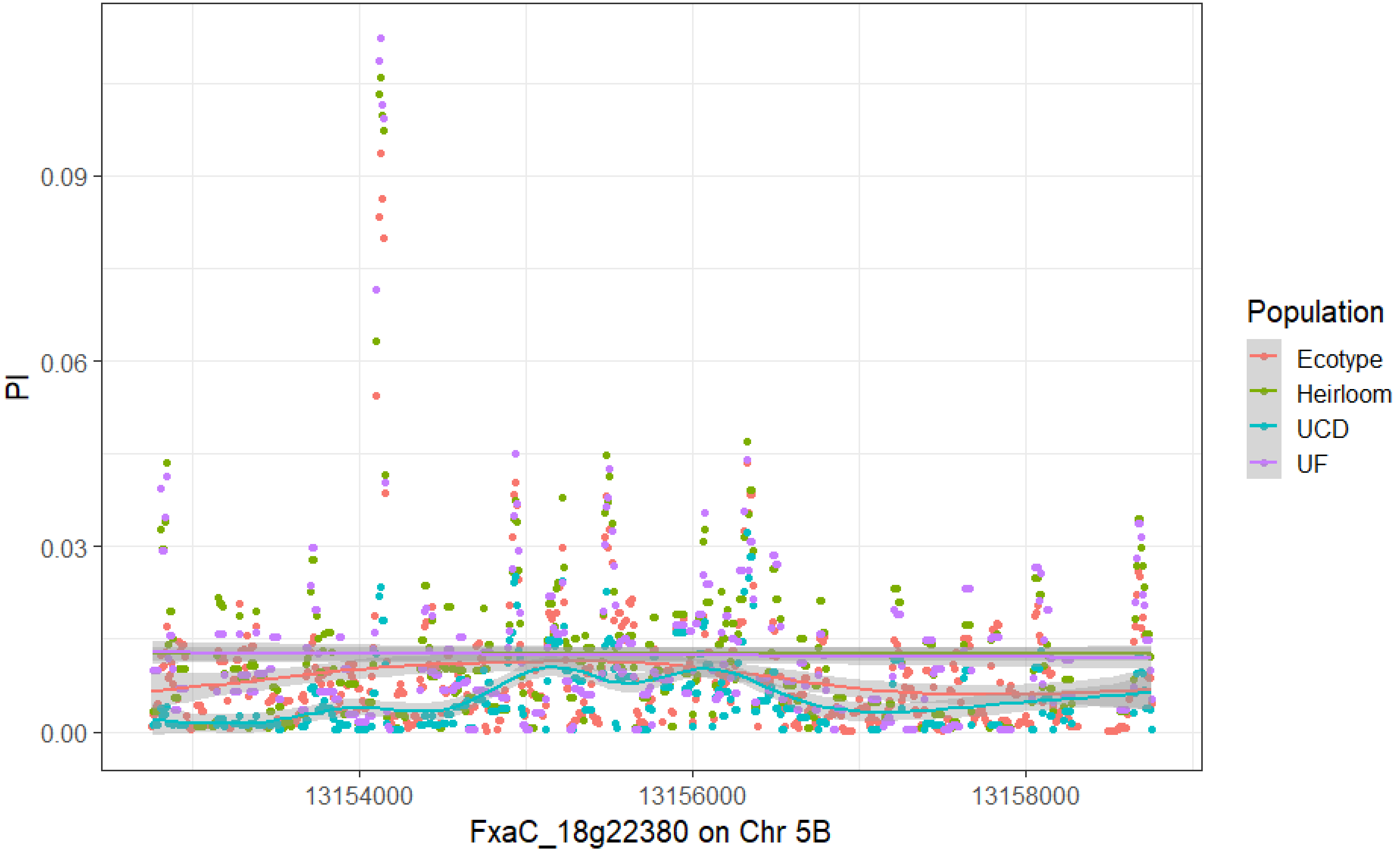


Figure S15. Nucleotide diversity (π) was measure in windows of 50 sites with a step size of 10. Lines are “GAM” smoothed distribution of π over the genic region of FxaC_18g22380 for each population.

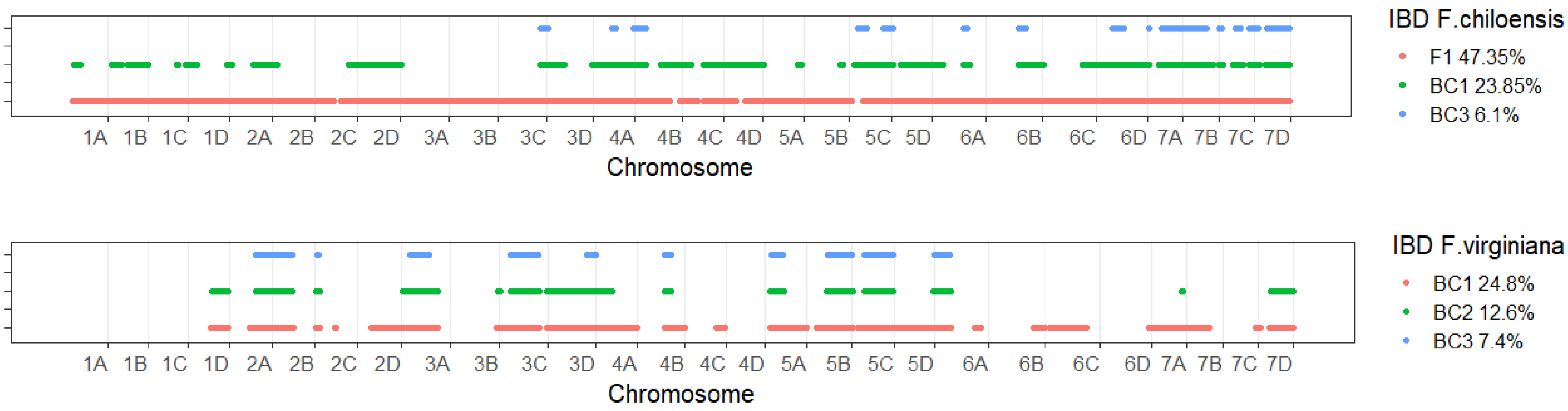


Figure S16. Identity by descent (IBD) segments in multiple generations of two backcrosses. Each generation is color coded. The genome wide IBD percentage in each generation is displayed in the legend box.

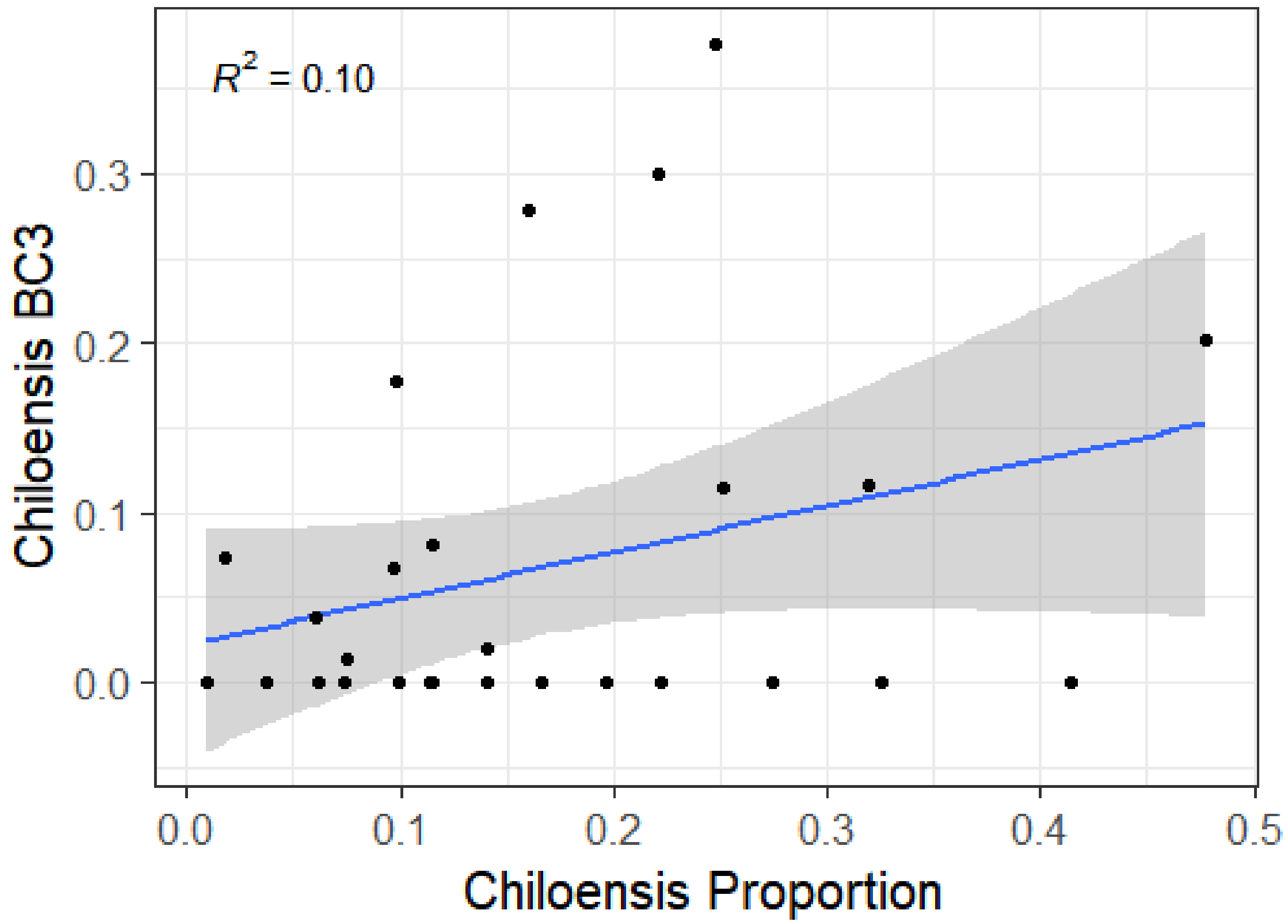


Figure S17. Chromosomal IBD percentage of the *Fragaria chiloensis* BC3 regressed on chromosomal proportion of *F. chiloensis* ancestry in the UF population.

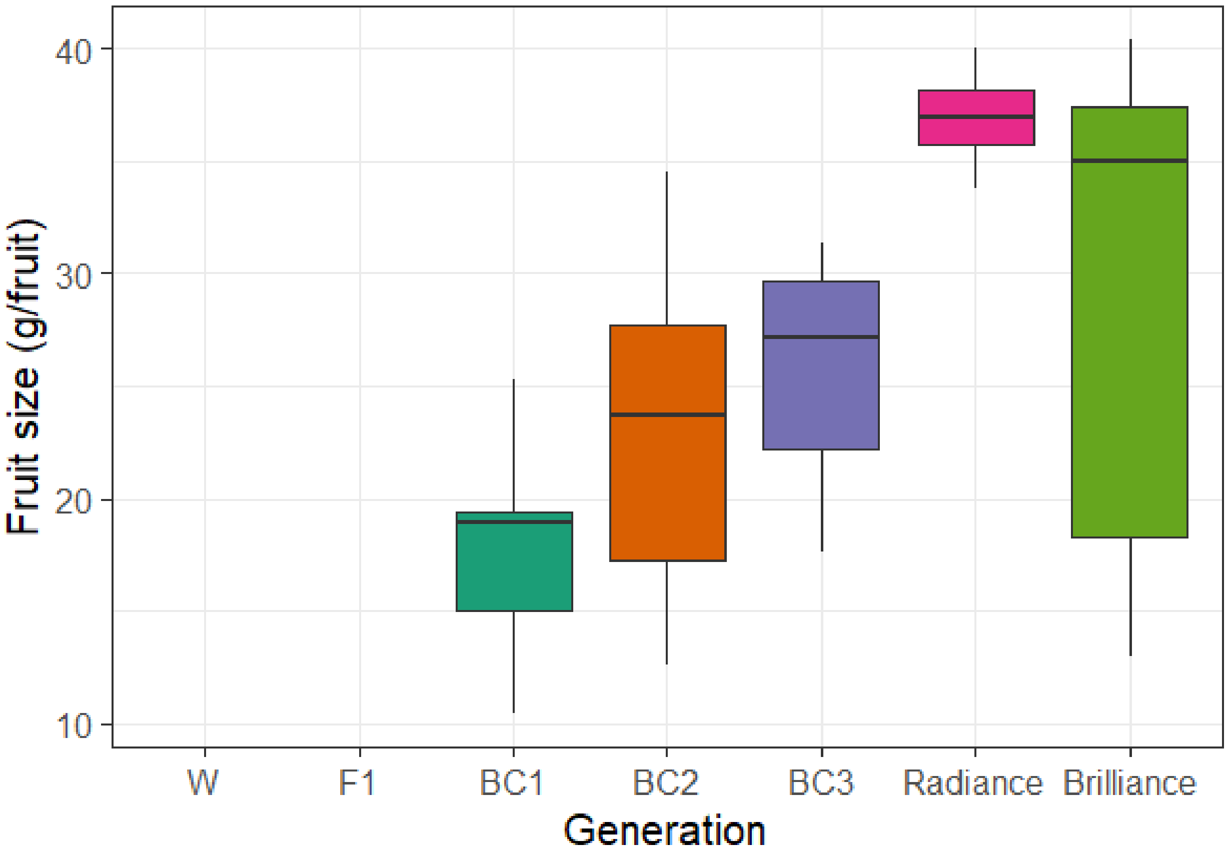


Figure S18. Comparisons of average fruit size among different backcross generations and two contemporary UF cultivars. W represents wild parents. Both W and F1 had no fruits across seven harvests, thus no data was recorded.

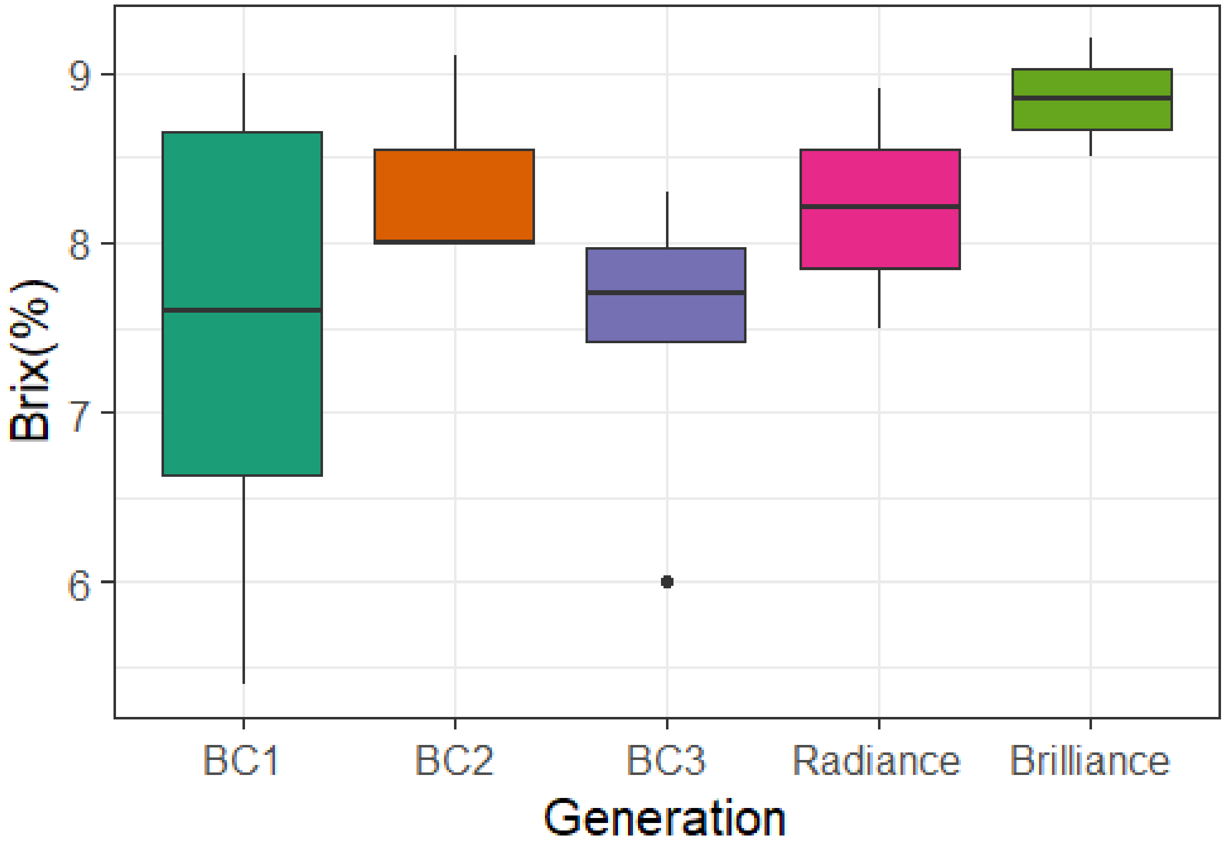


Figure S19. Comparisons of soluble solids content (Brix, %) among different backcross generations and two contemporary UF cultivars.
